# FBP1 is a nonenzymatic safety valve that curtails AKT activation to prevent insulin hyperresponsiveness

**DOI:** 10.1101/2023.03.05.531181

**Authors:** Li Gu, Yahui Zhu, Kosuke Watari, Maiya Lee, Junlai Liu, Sofia Perez, Melinda Thai, Joshua Mayfield, Bichen Zhang, Karina Cunha e Rocha, Fuming Li, Laura C. Kim, Alexander C. Jones, Igor H Wierzbicki, Xiao Liu, Alexandra C. Newton, Tatiana Kisseleva, Jun Hee Lee, Wei Ying, David J. Gonzalez, Alan R. Saltiel, M. Celeste Simon, Michael Karin

## Abstract

Insulin inhibits gluconeogenesis and stimulates glucose conversion to glycogen and lipids. How these activities are coordinated to prevent hypoglycemia and hepatosteatosis is not clear. Fructose-1,6 bisphosphatase (FBP1) is rate controlling for gluconeogenesis. However, inborn human FBP1 deficiency does not cause hypoglycemia unless accompanied by fasting or starvation, which also trigger paradoxical hepatomegaly, hepatosteatosis, and hyperlipidemia in affected individuals. Hepatocyte FBP1-ablated mice exhibit identical fasting-conditional pathologies along with AKT hyperactivation, whose inhibition reversed hepatomegaly, hepatosteatosis and hyperlipidemia but not hypoglycemia. Surprisingly, fasting-mediated AKT hyperactivation is insulin-dependent. FBP1 prevents insulin hyperresponsiveness, independently of its catalytic activity, by interacting with AKT, PP2A-C and Aldolase-B (ALDOB) to specifically accelerate AKT dephosphorylation. Enhanced by fasting and weakened by elevated insulin, FBP1:AKT:PP2A-C:ALDOB complex formation, which is disrupted by human FBP1 deficiency mutations or a C-terminal FBP1 truncation, prevents insulin-triggered liver pathologies and maintains lipid and glucose homeostasis. Conversely, a complex disrupting peptide reverses diet-induced insulin resistance.

## INTRODUCTION

Insulin is the master regulator of glucose homeostasis, keeping its blood concentration within a narrow range during fasting and feeding (Saltiel, 2021). A major site of insulin action is the liver, where it stimulates glucose uptake, glycolysis, and glucose conversion to fatty acids (FA) and glycogen, while inhibiting glucose output by blocking glycogenolysis (GL) and gluconeogenesis (GNG) (Hatting et al., 2018; Petersen et al., 2017). How these activities are integrated and regulated to prevent lethal hypoglycemia and avoid hepatosteatosis and hyperlipidemia, metabolic disorders that affect 30% of American adults, is not entirely clear. The key effector that controls glucose and lipid metabolism downstream to insulin during the fed state is the AKT-phosphoinositide-3-kinase (PI3K) pathway (Hay, 2011; Santoleri and Titchenell, 2019), but whether and how AKT is specifically deactivated during fasting is unknown.

While not a direct target for insulin signaling, fructose (F) 1,6 bisphosphatase 1 (FBP1) is rate controlling for GNG, converting F1,6-P_2_ to F6-P which is then converted to glucose (G) 6-P by phosphoglucoseisomerase (PGI) (Hunter et al., 2018; Li et al., 2014). FBP1 deficiency is a rare inborn metabolic error (OMIM:229700), that causes severe hypoglycemia, lactic acidosis, seizures, hepatomegaly, hyperlipidemia, hepatosteatosis and liver damage in carbohydrate starved, but not fed, infants (Gorce et al., 2022; Moey et al., 2018; Salih et al., 2020). While hypoglycemia and lactic acidosis are probably caused by low glycogen stores and abrogated GNG, there is no simple biochemical explanation for the hepatomegaly and elevated hepatic lipogenesis, which also affect non-starved older FBP1-deficient individuals (Gorce et al., 2022). We investigated hepatocyte specific *Fbp1* knockout mice (*Fbp1^ΔHep^*), initially generated for understanding FBP1’s tumor suppressive function in hepatocellular carcinoma (HCC) (Li et al., 2020), and found that fasted *Fbp1^ΔHep^* mice were phenotypically and metabolically identical to starved FBP1-deficient infants. In addition to low glycogen stores and severe hypoglycemia, these mice manifest fasting-induced liver pathologies, including hepatomegaly, hepatosteatosis and hyperlipidemia. Unexpectedly and independently of its catalytic activity, we discovered that FBP1 is a critical regulator of insulin signaling, serving as an endogenous “safety valve” that prevents insulin hyperresponsiveness and balances glucose and lipid metabolism. FBP1 negatively controls insulin signaling by nucleating a multiprotein complex that also contains Aldolase B (ALDOB), an enzyme that acts upstream to FBP1 in the GNG pathway (Figure S1A), the catalytic subunit of protein phosphatase 2A (PP2A-C), which dephosphorylates AKT (Schultze et al., 2012), and AKT, the key effector of insulin signaling (Hay, 2011). Complex formation that keeps AKT activation in check is enhanced by fasting and weakened at the fed state by insulin. Defective complex assembly results in insulin hyperresponsiveness, hepatosteatosis and hyperlipidemia in response to fasting-related insulin release by lipolysis generated free fatty acids (FFA) (Grill and Qvigstad, 2000) while preventing glucose intolerance. Conversely, pharmacological complex disruption reversed diet-induced insulin resistance.

## RESULTS

### Hypoglycemia, hepatomegaly, fatty liver, and hyperlipidemia in fasted *Fbp1^ΔHep^* mice

To understand the metabolic impact of FBP1 deficiency and its biochemical sequalae we crossed *Fbp1^F/F^* and *Alb-Cre* mice to generate *Fbp1^ΔHep^* mice. At 8 weeks-of-age, *Fbp1^ΔHep^*, but not *Fbp1^F/F^,* mice exhibited hepatomegaly, hepatosteatosis, severe hypoglycemia, hyperlipidemia, and signs of liver damage (elevated AST:ALT ratio) after a 16 h overnight fast (Figures 1A-1I). Weights of other tissues were not altered (Figure S1B). Hepatomegaly was due to hypertrophy rather than hyperplasia (Figures S1C and S1D). Elevated lipolysis was unlikely to be the cause of hepatosteatosis because adipose tissue mRNAs encoding hormone sensitive lipase (HSL) and adipose triglyceride lipase (ATGL) were insignificantly affected by *Fbp1* ablation (Figure S1E). Despite hypoglycemia, the *Fbp1^ΔHep^* livers contained more NADPH, ATP, and acetyl-CoA, which support lipid synthesis, than *Fbp1^F/F^* livers (Figures 1J, S1F-S1H). No genotype-related differences in serum insulin or glucagon were observed in the fast state, although serum lactate was elevated in fasted *Fbp1^ΔHep^* mice (Figures S1I-S1K). Consistent with Periodic Acid Schiff (PAS) staining, hepatic glycogen, and G6-P were reduced (Figures 1B, S1L and S1M). Interestingly, even after a mere 4 h fast, *Fbp1^ΔHep^* mice showed mild hepatomegaly, hepatosteatosis, hypoglycemia, hyperlipidemia, signs of liver injury and elevated hepatic NADPH:NADP ratio and ATP (Figures 1K-1T, S1N and S1O). No significant differences in weights of other tissues or lipolytic enzyme mRNAs were observed (Figures S1P and S1Q). Immunoblot (IB) analysis confirmed insignificant differences in HSL and ATGL phosphorylation in epididymal white adipose tissue (eWAT) after the 4 h fast (Figure S1R), ruling out enhanced lipolysis as the basis for the observed metabolic abnormalities. The short 4 h fast still led to elevated serum lactate in *Fbp1^ΔHep^* mice but little genotype-related differences in serum insulin and glucagon (Figures S1S-S1U). Lastly, hepatic glycogen and G6-P were significantly lower in *Fbp1^ΔHep^* mice than in *Fbp1^F/F^* mice (Figures 1L, S1V and S1W). In short, the metabolic phenotypes of hepatic steatosis, hypoglycemia, hepatomegaly, and hyperlipidemia associated with FBP1 deficiency appeared quickly within 4 h of fasting.

**Figure 1.**
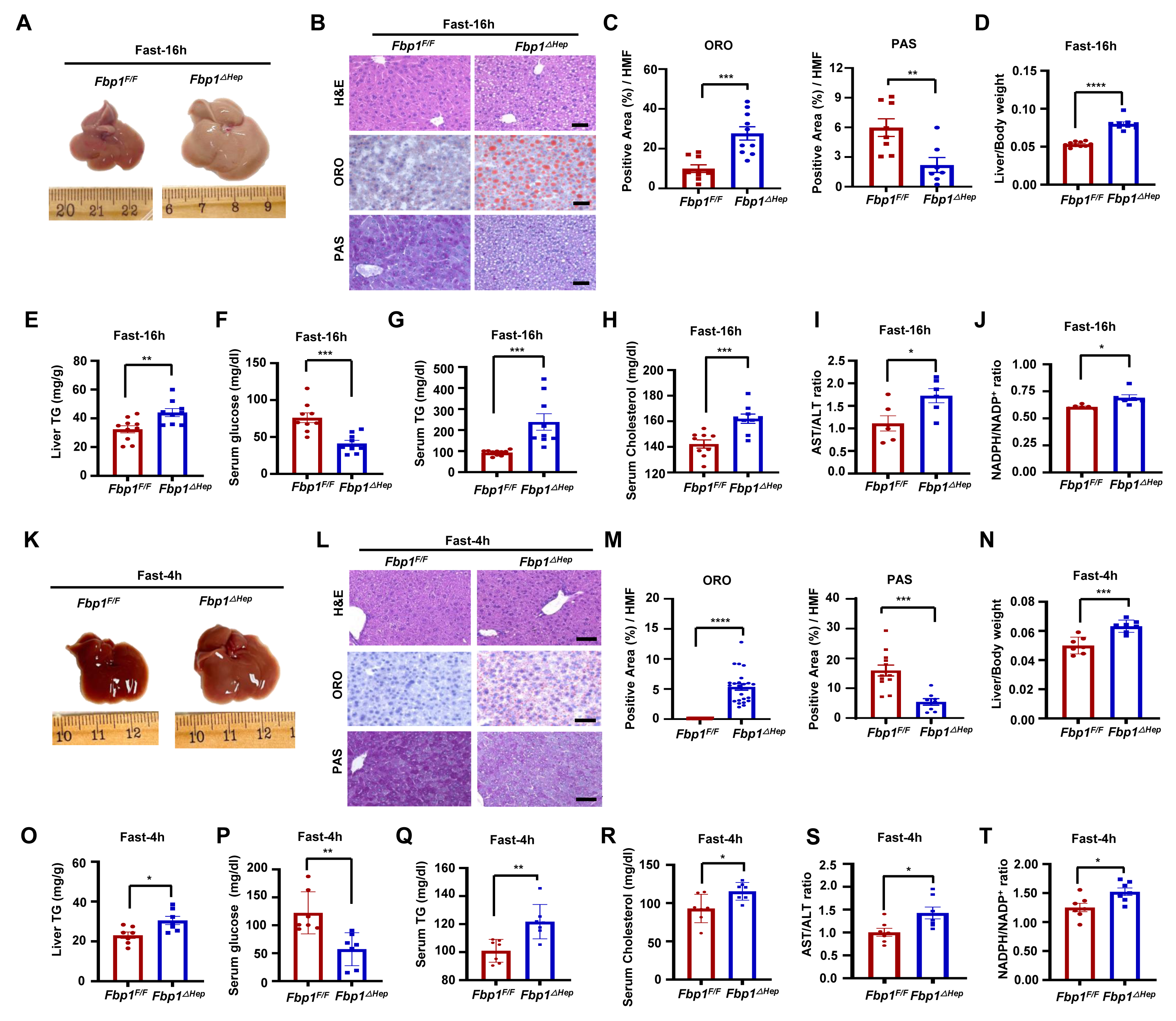
Hepatomegaly, hepatosteatosis, hypoglycemia in fasted *Fbp1^ΔHep^* mice. (A) Gross morphology of livers from 8-weeks-old (wo) *Fbp1^F/F^* and *Fbp1* ***^Δ^****^Hep^* mice fasted for 16 h. (B) Hematoxylin and eosin (H&E), Oil Red O (ORO), and periodic acid Schiff (PAS) staining of liver sections from 1A (n=8). scale bars, 20 μm. (C) ORO and PAS staining intensity per high-magnification-field (HMF) was determined by Image J from the slides shown in 1B. (D-H) Liver/Body weight (D), liver triglycerides (TG) (E), and serum glucose (F), TG (G), and cholesterol (H) in the indicated mice (n=9-10). (I and J) Serum aspartate aminotransferase (AST) / alanine aminotransferase (ALT) ratio (I) and liver NADPH/NADP^+^ ratio (J) in the indicated mice (n=6). (K) Gross morphology of livers from 8-weeks-old (wo) *Fbp1^F/F^* and *Fbp1^ΔHep^* mice fasted for 4 h. (L) H&E, ORO, and PAS staining of liver sections from K (n=7). scale bars, 20 μm. (M) ORO and PAS staining intensity per HMF was determined by Image J analysis of the data in 1L. (N-R) Liver/Body weight (N), liver TG (O), and serum glucose (P), TG (Q), and cholesterol (R) in the indicated mice (n=7). (S and T) Serum AST/ALT ratio (S) and liver NADPH/NADP^+^ ratio (T) in the indicated mice (n=7). Data are presented as mean ± SEM. *P < 0.05, **P < 0.01, ***P < 0.001 ****p < 0.0001 (Unpaired two-tailed t test).

### Fasted *Fbp1^ΔHep^* mice exhibit enhanced AKT-mTOR signaling and de novo lipogenesis

Fasted (16 h) *Fbp1^ΔHep^* livers and primary mouse hepatocytes showed enhanced AKT S473 and T308 and GSK3β S9 phosphorylation, loss of AMPK T172 phosphorylation and elevated S6 and p70 S6 kinase (p70S6K) phosphorylation, suggesting mTORC1 activation (Figures 2A and S2A), Consistent with hepatosteatosis, previous findings (Li et al., 2020), and mTORC1 and p70S6K activation (Bakan and Laplante, 2012), nuclear/mature (m) SREBP1c was strongly elevated in fasted *Fbp1^ΔHep^* livers, along with its targets, ATP citrate lyase (ACLY), acetyl CoA carboxylase 1 (ACC1) and fatty acid synthase (FASN) and the SREBP2 targets HMG CoA synthase 1 (HMGCS1) and reductase (HMGCR) (Figures 2B-2D). The lipid transporter CD36, Glut2, pyruvate kinase M2 (PKM2), phosphogluconate dehydrogenase (PGD), transketolase (TKT), malic enzyme 1 (ME1), methylenetetrahydrofolate dehydrogenase 2 (MTHFD2), phosphoribosyl pyrophosphate amidotransferase (PPAT) and 1,4-α-glucan branching enzyme 1 (GBE1) mRNAs were elevated, whereas carnitine palmitoyl transferases 1a and 2 (CPT1a, CPT2) mRNAs, involved in β-oxidation, were reduced (Figures 2E-2G). Carbohydrate response element binding protein (ChREBP), whose overexpression promotes hepatic steatosis while improving insulin sensitivity (Benhamed et al., 2012), was also elevated in the cytoplasm and nucleus of *Fbp1^ΔHep^* livers and so was glucocorticoid receptor (GR) (Figure S2B).

**Figure 2.**
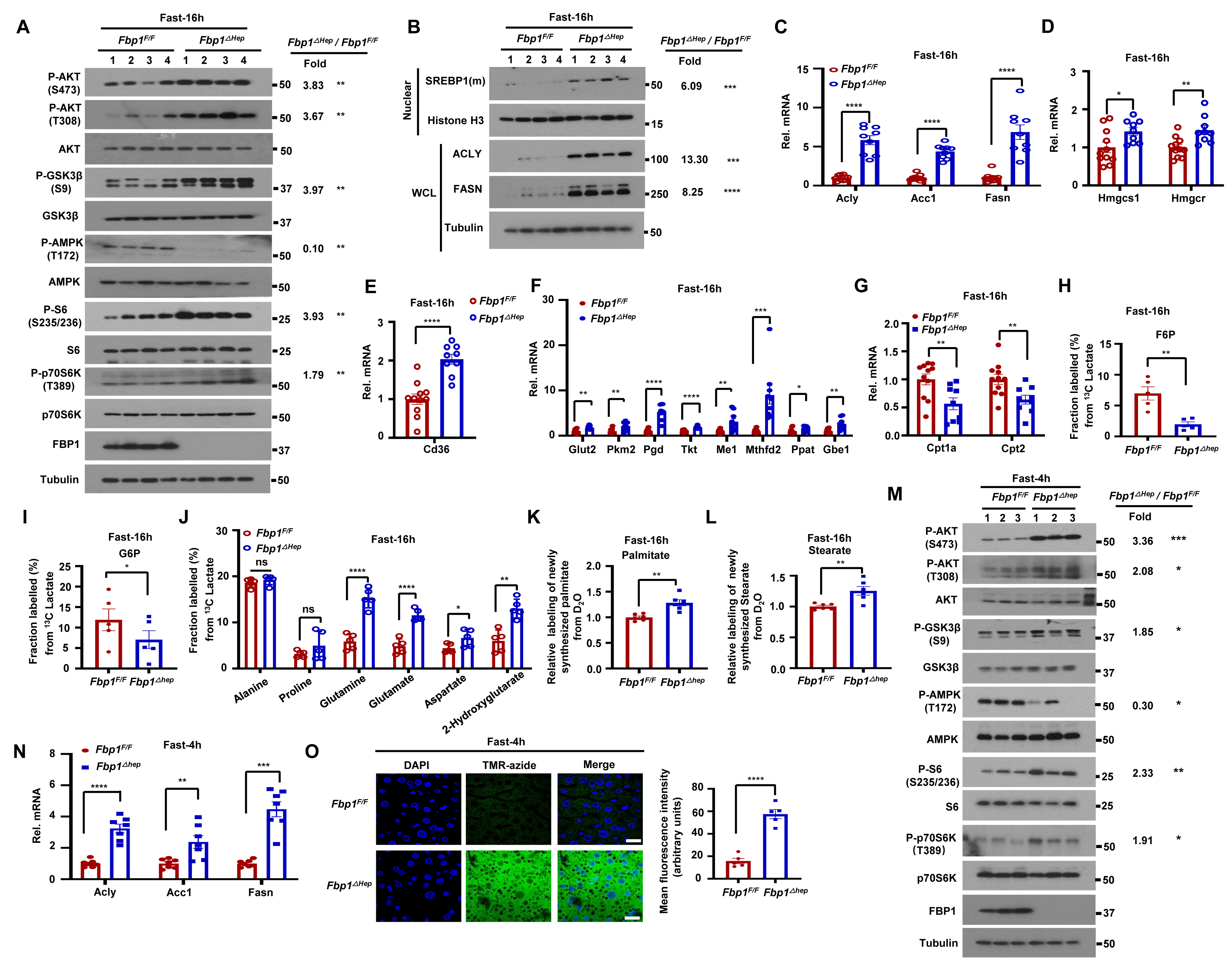
Fasted *Fbp1^ΔHep^* mice exhibit enhanced AKT-mTOR signaling and elevated de novo lipogenesis. (A) Immunoblot (IB) analysis of the indicated proteins in livers of 16 h fasted 8-wo *Fbp1^F/F^* and *Fbp1^ΔHep^* mice. Densitometric quantification of relative normalized protein ratios (*Fbp1^ΔHep^/Fbp1^F/F^*) and P values are shown on the right. (B) IB analysis of indicated nuclear and cytoplasmic proteins in 16 h fasted 8-wo *Fbp1^F/F^* and *Fbp1^ΔHep^* mice (n=4). Relative normalized protein ratios (*Fbp1^ΔHep^/Fbp1^F/F^*) and P values are shown on the right. (C-G) qRT-PCR analysis of lipogenesis (C), cholesterol synthesis (D), lipid uptake (E), glycolysis, pentose phosphate pathway, glycogen synthesis (F) and β-oxidation (G) related liver mRNAs from fasted 8-wo *Fbp1^F/F^* and *Fbp1^ΔHep^* mice (n=9-10) (H-J) Fractional labeling of liver F6-P (H), G6-P (I) and amino acids (J) from ^13^C-lactate tracing of 16 h fasted 8-wo *Fbp1^F/F^* and *Fbp1^ΔHep^* mice (n=5). (K and L) Relative deuterium incorporation into newly synthesized liver palmitate (C16:0) (K) and stearate (C18:0) (L) after 24 h of D2O labeling (n=4-6). (M) IB analysis of indicated liver proteins from 4 h fasted 8-wo *Fbp1^F/F^* and *Fbp1^ΔHep^* mice. Relative normalized protein ratios (*Fbp1^ΔHep^/Fbp1^F/F^*) and P values are shown on the right. (N) qRT-PCR analysis of lipogenesis related liver mRNAs from 4 h fasted 8-wo *Fbp1^F/F^* and *Fbp1^ΔHep^* mice (n=7). (O) Liver protein synthesis in 4 h fasted 8-wo *Fbp1^F/F^* and *Fbp1^ΔHep^* mice was imaged (left) by O-propargyl-puromycin (OPP, TMR-azide) incorporation (n=5), and mean fluorescence intensity was quantified (right). Scale bars, 20 μm. Data are presented as mean ± SEM. *P < 0.05, **P < 0.01, ***P < 0.001 ****p < 0.0001 (Unpaired two-tailed t test). All blots are representative of at least 3 biological replicates.

GC-MS based metabolomic analysis of ^13^C-glucose loaded livers revealed elevated glycolytic flux and decreased TCA cycle activity in fasted *Fbp1^ΔHep^* livers (Figures S2C-S2E), explaining the increase in serum lactate. ^13^C-lactate tracing showed reduced lactate flux to F6-P and G6-P but enhanced lactate flux into the TCA cycle and amino acids (Figures 2H-2J, S2C and S2F), correlating with reduced liver glycogen. D_2_O tracing confirmed elevated de novo lipogenesis (DNL) in fasted *Fbp1^ΔHep^* mice (Figures 2K and 2L). Curiously, glycogen synthase 2 (GYS2) and its phosphorylated form (P-GS) were elevated in fasted *Fbp1^ΔHep^* livers (Figure S2G), despite their low glycogen content and the increase in inhibitory GSK3 phosphorylation, which should have increased GYS2 activity (Beurel et al., 2015). This suggested that low liver glycogen stores in these mice could be due to low G6-P, the key allosteric activator of GYS2 (Villar-Palasí and Guinovart, 1997), which is generated by PGI from F6-P, whose production from lactate via GNG was reduced. Indeed, GYS activity was significantly lower in *Fbp1^ΔHep^* liver lysates relative to *Fbp1^F/F^* liver lysates (Figure S2H). This difference was indeed due to lower G6-P in the *Fbp1^ΔHep^* liver because when exogenous G6-P was added to the reaction mix, GYS activity was higher in *Fbp1^ΔHep^* liver lysates, consistent with its higher abundance.

After a 4 h fast, *Fbp1^ΔHep^* livers also showed elevated AKT activation, GSK3β S9 phosphorylation, S6 and p70S6K phosphorylation and reduced AMPK activation (Figure 2M). SREBP1 target mRNAs encoding ACLY, ACC1 and FASN were moderately elevated, along with the SREBP2 target HMGCR, the lipid transporter CD36, and the metabolic enzymes PKM2, PGD, TKT, ME1 and PPAT, whereas CPT2 mRNA was decreased (Figures 2N and S2I-S2L). Consistent with mTORC1 activation and hepatomegaly, protein synthesis measured with O-propargyl-puromycin (OP-puro) was strongly elevated in 4 h-fasted *Fbp1^ΔHep^* hepatocytes (Figure 2O) and so were nuclear ChREBP and GR (Figure S2M). Thus, the FBP1 deficiency results in rapid fasting induced upregulation of AKT-mTOR signaling, lipogenesis and protein synthesis.

Of note, in the fed state *Fbp1^ΔHep^* livers were barely different from *Fbp1^F/F^* livers, although the *Fbp1^ΔHep^* liver exhibited slight hepatomegaly, somewhat lower glycogen and marginally elevated liver damage (Figures S3A-S3J). Serum parameters also did not differ between the two genotypes (Figures S3K-S3O). *Fbp1^ΔHep^* livers also showed increased ACLY and ME1 mRNAs, but mRNAs coding for other metabolic genes did not differ from those in *Fbp1^F/F^* livers (Figures S3P-S3T). No changes in GYS2 and P-GS expression were observed (Figure S3U).

### AKTi and mTORCi attenuate lipogenesis and protein synthesis but not hypoglycemia

To determine whether elevated AKT or mTORC1 activities contribute to the metabolic abnormalities in fasted *Fbp1^ΔHep^* mice, we treated the mice with AKT (MK2206) or mTORC1/2 (Torin1) inhibitors 28 and 4 h before fasting. Both inhibitors prevented hepatosteatosis and hypertriglyceridemia, reducing them to levels seen in drug-treated fasted *Fbp1^F/F^* mice, but had no effect on hypoglycemia (Figures 3A-3G). Serum cholesterol was reduced by MK2206 but not by Torin1 (Figure 3H). The two inhibitors partially abrogated hepatomegaly (Figures 3A and 3I) and blunted the fasting-induced increase in protein synthesis but did not alter serum insulin (Figures S4A-S4C). As expected, MK2206 and Torin1 (which also inhibits mTORC2 that acts upstream to AKT) blocked fasting induced AKT, ribosomal protein S6 and GSK3β phosphorylation and blunted SREBP1c, ACLY, FASN, HMGCR and CD36 mRNA expression (Figures S4D and S4E).

**Figure 3.**
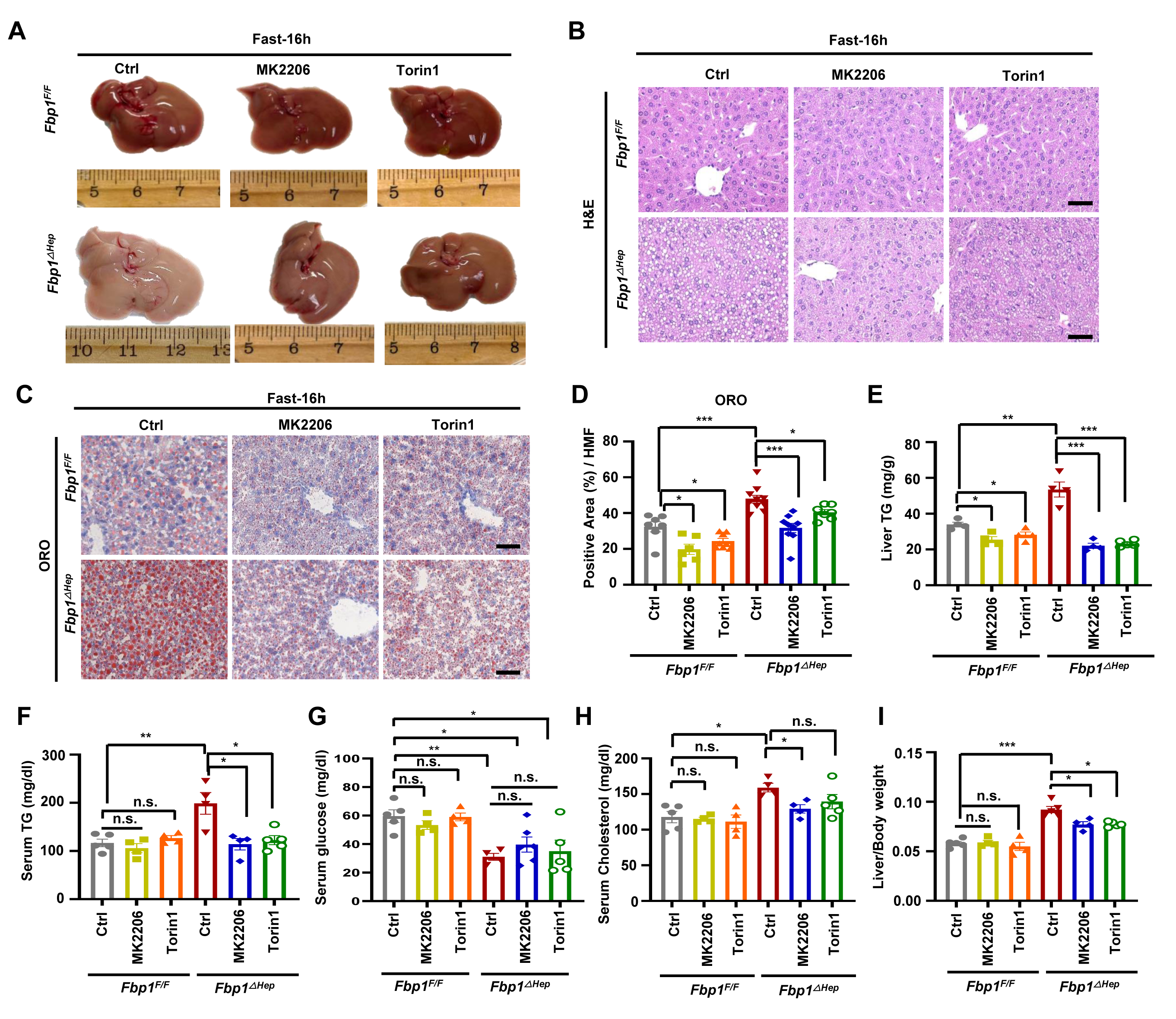
AKT inhibition attenuate hepatomegaly, hepatosteatosis and hyperlipidemia but not hypoglycemia. (A) Gross liver morphology of 8-wo *Fbp1^F/F^* and *Fbp1^ΔHep^* mice treated -/+ MK2206 (100 mg/kg) or Torin1 (10 mg/kg), followed by 16 h fast before sacrifice. (B-D) H&E (B) and ORO staining (C) of liver sections from 3A (n=4-5). Scale bars, 20 μm. ORO staining intensity per HMF was determined by Image J (D). (E-I) Liver TG (E), serum TG (F), serum glucose (G), serum cholesterol (H) and liver/body weight ratio (I) in indicated mice (n=4-5). Data are presented as mean ± SEM. n.s., not significant, P≥0.05. *P < 0.05, **P < 0.01, ***P < 0.001 (Unpaired two-tailed t test).

### Catalytically inactive FBP1 prevents hepatomegaly, steatosis, and hyperlipidemia

In addition to catalyzing GNG, FBP1 non-enzymatically inhibits HIF-1 activity (Li et al., 2014). AAV8-mediated re-expression of WT FBP1 or catalytically inactive FBP1^E98A^ in *Fbp1^ΔHep^* hepatocytes had an effect akin to AKTi treatment, blunting fasting-induced hepatomegaly, hepatosteatosis and hyperlipidemia and reducing lipogenic gene expression, although as expected FBP1^E98A^ did not obliterate hypoglycemia (Figures 4A-4H, S5A and S5B). AAV8 mediated FBP1 or FBP1^E98A^ re-expression largely suppressed AKT activation and GSK3β S9 phosphorylation (Figures S5C and S5D).

**Figure 4.**
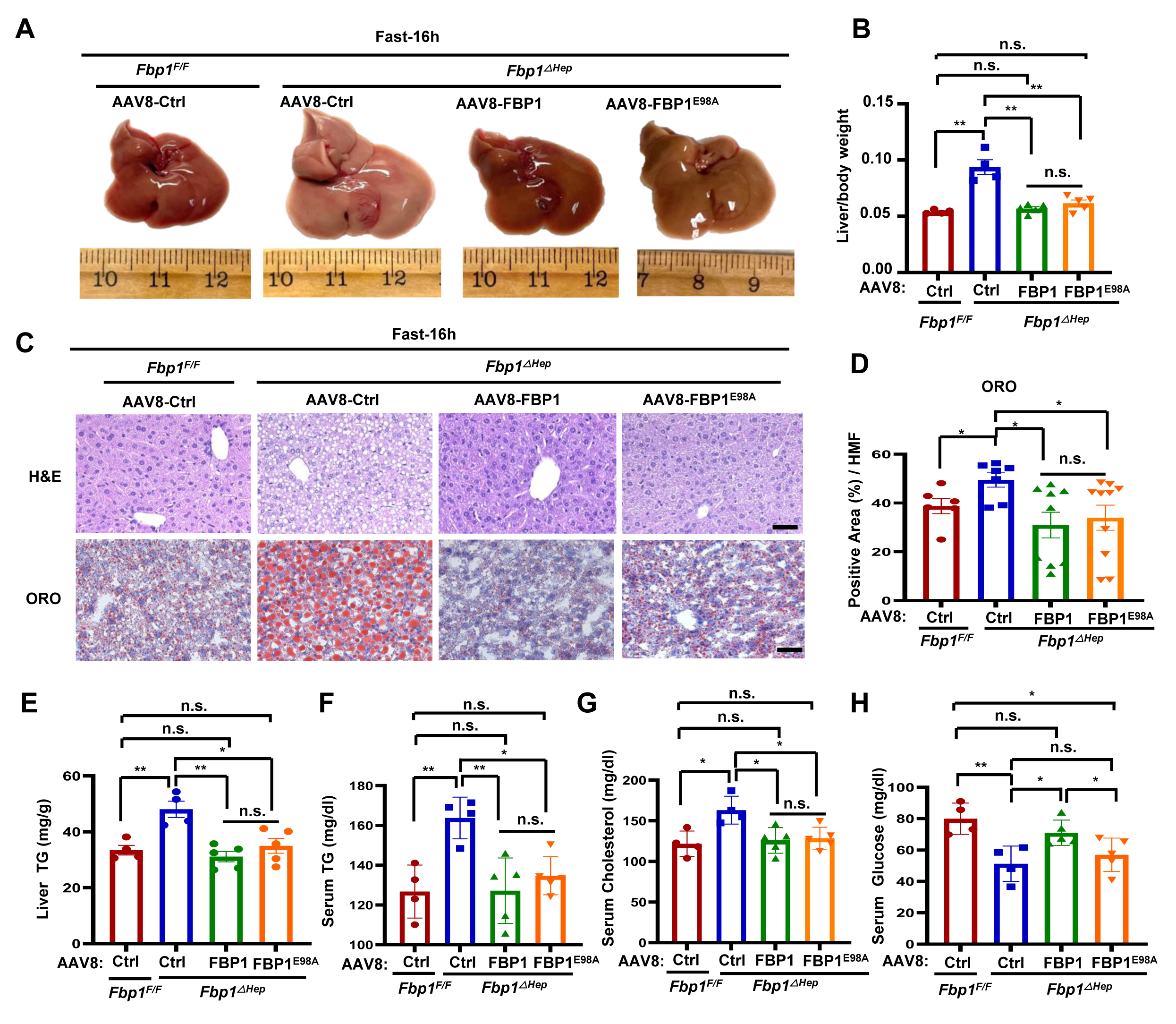
Inactive FBP1 prevents hepatomegaly, steatosis, and hyperlipidemia. (A) Gross liver morphology of 8-wo *Fbp1^F/F^* and *Fbp1^ΔHep^* transduced with AAV8-Ctrl, AAV8-FBP1 and AAV8-FBP1^E98A^, followed by 16 h fast before sacrifice at 3 wk post-AAV8 infection (n=4-5). (B) Liver/body weight ratio in above mice (n=4-5). (C and D) H&E and ORO staining of liver sections from 4A (n=4-5) (C). Scale bars, 20 μm. ORO staining intensity per HMF determined by Image J (D). (E-H) liver TG (E), serum TG (F), serum cholesterol (G), and serum glucose (H) in the indicated mice (n=4-5). Data are mean ± SEM. n.s., not significant, P≥0.05, *P < 0.05, **P < 0.01 (Unpaired two-tailed t test).

To elucidate how FBP1 regulated AKT activation, we used mass spectrometry (MS) as an unbiased tool for the identification of FBP1 binding proteins. ALDOB, AKT1 and PP2A-C ranked as the top FBP1 interacting partners present in HA-FBP1 immunoprecipitates (IP’s) isolated from AAV8-FBP1 transduced *Fbp1^ΔHep^* livers (Figure S5E). *Fbp1^ΔHep^* livers transduced with AAV8-Ctrl served as a negative control. Of note, no PP2A regulatory subunits co-IP’ed with FBP1.

### FBP1 associates with PP2A-C and ALDOB to interact with AKT and inhibit its activation

Next, we examined whether PP2A-C and ALDOB were part of the AKT inhibitory mechanism triggered by FBP1. FBP1-deficient individuals exhibit mild hereditary fructose intolerance (HFI) (Gorce et al., 2022; Moey et al., 2018), a metabolic disorder associated with ALDOB deficiency (OMIM;229600), which blocks F1,6-P_2_ breakdown (Herman and Birnbaum, 2021; Simons et al., 2019). ALDOB deficiency also causes hepatomegaly, hepatosteatosis and liver damage (Di Dato et al., 2019; Oppelt et al., 2015). This phenotypic overlap suggested that FBP1 may regulate AKT activity through its interaction with ALDOB, which was also detected in previous studies (Droppelmann et al., 2015). Consistent with our MS analysis, ALDOB was reported to recruit PP2A-C to AKT in HCC cells and enhance AKT dephosphorylation (He et al., 2020b). Indeed, we confirmed that ectopically expressed WT and catalytically inactive FBP1^E98A^ interacted with AKT1, AKT2, ALDOB and PP2A-C in 293T cells, and that ectopic ALDOB IPs contained endogenous AKT1 and FBP1 (Figures S6A and S6B). Likewise, ectopic PP2A-C interacted with AKT1 and FBP1, which enhanced the AKT1-PP2A-C interaction (Figure S6C). Ectopic WT or FBP1^E98A^ expression in Huh7 cells potentiated the interaction between AKT, ALDOB and PP2A-C, but FBP1 or ALDOB silencing disrupted the interactions between AKT, PP2A-C, ALDOB and FBP1 (Figures S6D-S6F). These interactions were also confirmed in mouse liver; AKT IPs from *Fbp1^F/F^* or *Fbp1^ΔHep^* livers reconstituted with FBP1 or FBP1^E98A^ contained FBP1, ALDOB and PP2A-C (Figure 5A). Similar results were obtained when AKT was IP’ed from primary mouse hepatocytes, in which the absence of FBP1 diminished the association of AKT with ALDOB and PP2A-C (Figure 5B), indicating that FBP1 plays a pivotal role in complex formation. Formation of the FBP1-dependent AKT:PP2A-C:ALDOB complex was enhanced by fasting (Figures 5C, S6G and S6H). In vitro binding experiments confirmed assembly of the FBP1:PP2A-C:ALDOB:AKT complex using purified recombinant proteins, revealing that FBP1 first binds ALDOB and PP2A-C, and weakly to AKT1 (Figure 5D-F). Addition of ALDOB and PP2A-C enhanced binding to AKT1 and although ALDOB and PP2A-C interacted with AKT1 these interactions are enhanced by FBP1. Gel filtration chromatography of *Fbp1^F/F^* liver lysates revealed co-elution of FBP1, AKT, PP2A-C and ALDOB in a high molecular weight fraction, averaging 750-800 kDa, which was not seen in *Fbp1^ΔHep^* lysates (Figure S6I and S6J).

**Figure 5.**
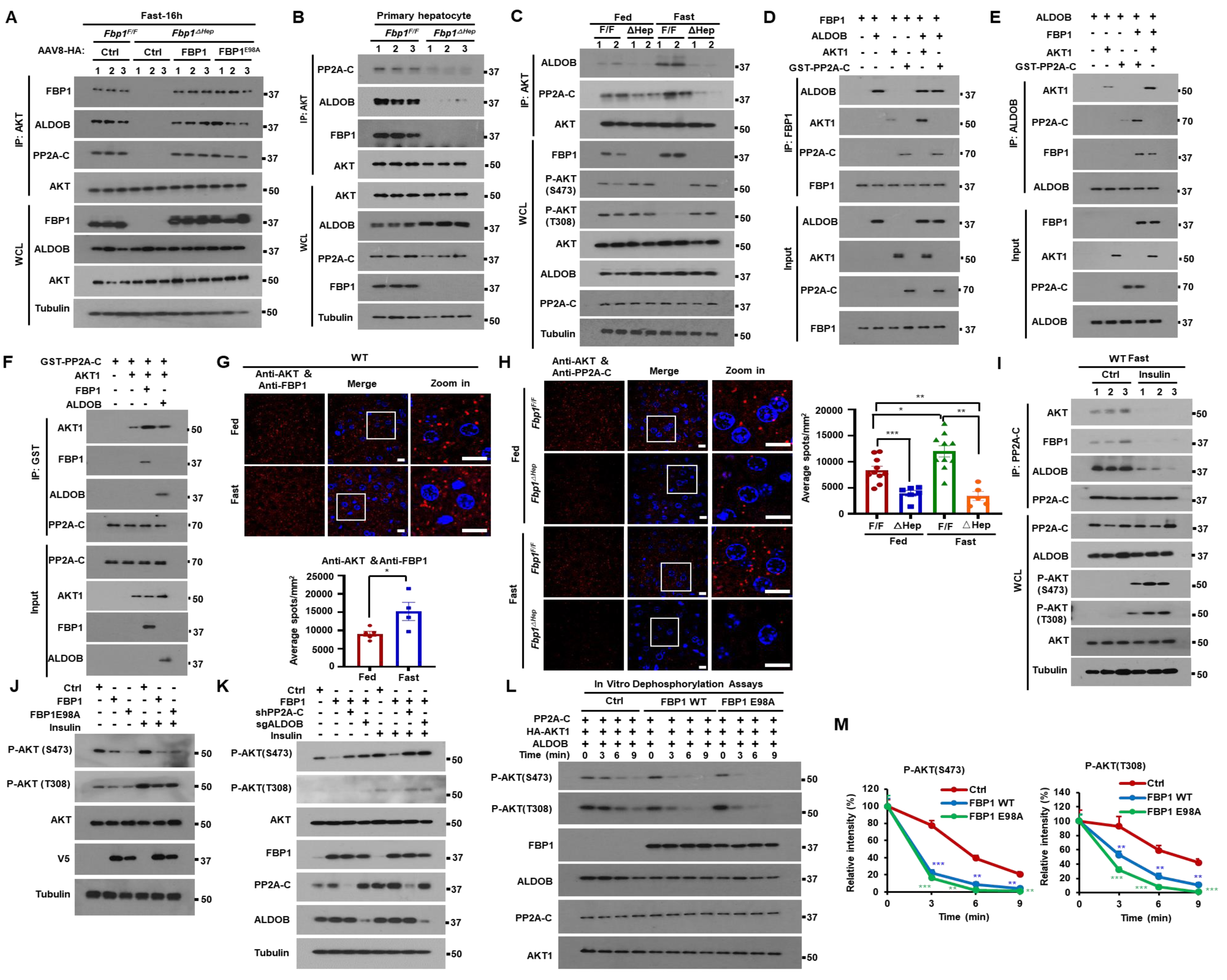
FBP1 associates with PP2A-C and ALDOB to inhibit AKT activation. (A) IP analysis of liver lysates from *Fbp1^F/F^* and *Fbp1^ΔHep^* mice transduced with AAV8-Ctrl, AAV8-FBP1 and AAV8-FBP1^E98A^ (n=3). The gel separated IPs were IB’ed with the indicated antibodies. (B) IP of primary mouse hepatocytes isolated from 16 h fasted *Fbp1^F/F^* and *Fbp1^ΔHep^* mice (n=3). The gel separated IPs were IB’ed with the indicated antibodies. (C) IP analysis of liver lysates from fed or 16 h fasted 8-wo *Fbp1^F/F^* and *Fbp1^ΔHep^* mice. The gel separated IPs were IB’ed with the indicated antibodies. (D-F) In vitro associations between FBP1 (D), ALDOB (E), and GST-PP2A-C (F) with the indicated recombinant proteins. The gel separated IPs were IB’ed with the indicated antibodies. (G and H) PLA of AKT-FBP1 (G, top) and AKT-PP2A-C (H, left) interactions in liver sections of fed or 16 h fasted 8-wo *Fbp1^F/F^* and *Fbp1^ΔHep^* mice (n=5-8). Scale bars, 10 μm. Quantification of representative stained tissues (G, bottom and H, right). (I) IP of liver lysates of WT mice treated -/+ insulin (0.5 U/kg) after an overnight fast (n=3). The gel separated IP’s were IB’ed with indicated antibodies. (J and K) IB analysis of Huh7 cells stably transfected with the indicated vectors and stimulated or not with insulin (100 nM) for 1 h after 6 h of serum starvation. (L and M) In vitro dephosphorylation of phosphorylated HA-AKT1 isolated from insulin stimulated Huh7 cells and incubated with active-PP2A-C in the presence or absence of indicated recombinant proteins. The reactions were IB analyzed with the indicated antibodies (L). Relative intensity of P-AKT (S473) and P-AKT (T308) was determined by densitometry (M). Data are presented as mean ± SEM. *P < 0.05, **P < 0.01, ***P < 0.0001 (Unpaired two-tailed t test).

Proximity ligation assays (PLA) confirmed formation of the FBP1:PP2A-C:ALDOB:AKT complex in mouse liver and the pivotal role of FBP1 in coordinating ALDOB and PP2A-C recruitment to AKT (Figures 5G-5H and S6K-S6P). The interactions between FBP1, ALDOB, PP2A-C and AKT were enhanced by fasting and weakened by feeding or insulin (Figures 5G-5I and S6K-S6V). FBP1 knockdown in the kidney epithelial cell line HK-2 and the colon carcinoma-derived CaCo-2 cell line also augmented AKT activation with and without insulin (Figures S6W). Ectopic expression of either WT FBP1 or FBP1^E98A^ in HCC cells inhibited basal and insulin-stimulated AKT S473 phosphorylation in a PP2A-C-and ALDOB-dependent manner (Figures 5J and 5K). In vitro dephosphorylation assay using active AKT1 as a substrate showed that either FBP1 isoform stimulated time-dependent S473 and T308 dephosphorylation in an ALDOB dependent manner (Figures 5L, 5M and S6X). Dephosphorylation of another PP2A substrate was not affected by FBP1 (Figure S6Y).

### *Fbp1^ΔHep^* mice are hyperresponsive to insulin

As obesity results in insulin resistance which interferes with AKT activation, we checked how HFD consumption affects the FBP1-PP2A-C-ALDOB-AKT complex. WB and PLA showed that HFD enhanced FBP1 and ALDOB expression as well as complex formation and inhibited AKT activation in WT mice (Figures S7A-S7D). We also fed *Fbp1^F/F^* and *Fbp1^ΔHep^* mice HFD for 12 weeks. While weight gain was nearly identical between the two, *Fbp1^ΔHep^* mice exhibited higher liver to body weight ratio (Figures S7E and S7F), hepatomegaly, enhanced hepatosteatosis but less liver glycogen than *Fbp1^F/F^* mice (Figures 6A-6C). Although HFD-fed *Fbp1^ΔHep^* mice were protected from obesity-related hyperglycemia, their liver and serum triglycerides (TG) and cholesterol were elevated (Figures 6D-6G). Glucose tolerance test (GTT) showed that HFD-fed *Fbp1^ΔHep^* mice were more glucose tolerant than *Fbp1^F/F^* mice, although insulin levels did not differ between the two (Figures 6H and S7G). Insulin tolerance test (ITT) confirmed insulin hypersensitivity in HFD-fed *Fbp1^ΔHep^* mice (Figure 6I), whose liver contained more phosphorylated AKT and GSK3β than *Fbp1^F/F^* liver (Figure 6J). Insulin administration to either HFD-or normal chow (NCD)-fed *Fbp1^F/F^* and *Fbp1^ΔHep^* mice resulted in higher AKT phosphorylation, under both HFD and NCD feeding and a considerable and sustained drop in blood glucose in NCD-fed *Fbp1^ΔHep^* mice (Figures 6K, 6L and S7H). GTT and ITT indicated that both FBP1 and FBP1^E98A^ re-expression decreased glucose tolerance and insulin hypersensitivity (Figures S7I and S7J), even though insulin levels during GTT did not differ between the two (Figures S7K and S7L). *Fbp1^ΔHep^* mice were less pyruvate tolerant and this was reversed by AAV-FBP1, but not AAV-FBP1^E98A^, re-expression (Figures S7M and S7N). Suppression of insulin-induced AKT activation was also seen in human hepatocytes, where FBP1 silencing enhanced insulin induced AKT activation in a dose-and time-dependent manner (Figures 6M and S7O).

**Figure 6.**
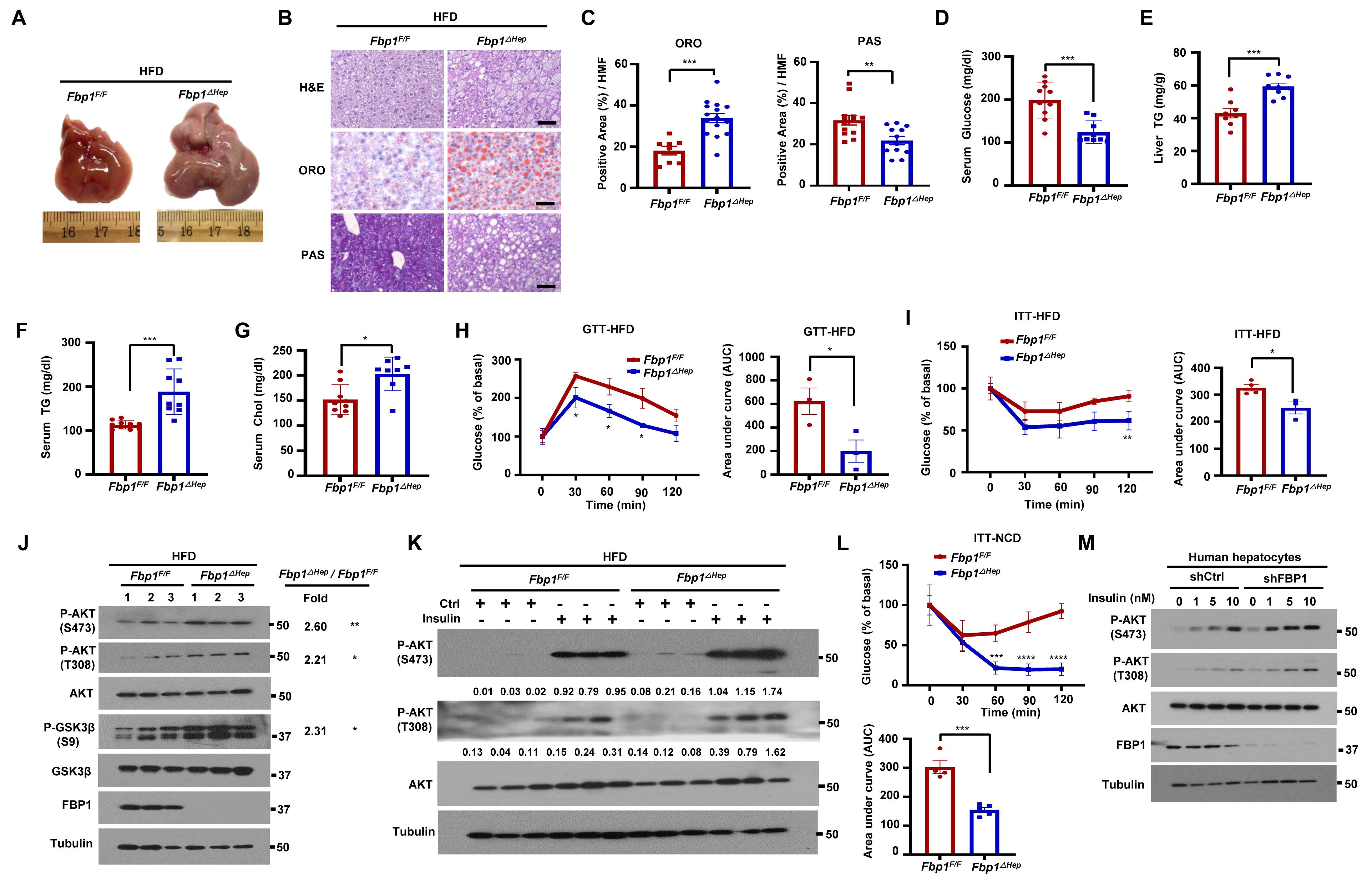
*Fbp1^ΔHep^* mice and FBP1-deficient human hepatocytes are insulin hyperresponsive. (A) Gross liver morphology of 18-wo HFD-fed *Fbp1^F/F^* and *Fbp1^ΔHep^* mice (n=8-10) that were fasted for 4 h before sacrifice. (B) H&E, ORO, and PAS staining of liver sections from above mice (n=8). Scale bars, 20 μm. (C) ORO and PAS staining intensity per HMF from 6C. (D-G) serum glucose (E), liver TG (F), serum TG (G) and serum cholesterol (H) in indicated HFD-fed mice (n=8-10). (H) Glucose tolerance test (GTT) of HFD-fed *Fbp1^F/F^* and *Fbp1^ΔHep^* mice (left) (n=3) and area under the curve (AUC) quantification (right). (I) Insulin tolerance test (ITT) of HFD-fed *Fbp1^F/F^* and *Fbp1^ΔHep^* mice (left) (n=4) and AUC quantification (right). (J) IB analysis of liver lysates from indicated HFD-fed mice. Densitometric quantification of relative normalized protein ratios (*Fbp1^ΔHep^/Fbp1^F/F^*) and P values are shown on the right. (K) IB analysis of liver lysates of HFD-fed 18-wo *Fbp1^F/F^* and *Fbp1^ΔHep^* mice collected 15 min after control or insulin injections (n=4). Quantification of relative normalized protein amounts is shown below each P-AKT strip. (L) ITT of NCD-fed *Fbp1^F/F^* and *Fbp1^ΔHep^* mice (top) (n=4-5) and AUC quantification (bottom). (M) Human hepatocytes stably transfected with shCtrl or shFBP1 were stimulated with the indicated concentrations of insulin for 1 h after 6 h of serum starvation were IB analyzed with the indicated antibodies. Data are mean ± SEM. *P < 0.05, **P < 0.01, ***P < 0.001 (Unpaired two-tailed t test).

Insulin concentrations are higher in the fed state relative to the fasted state, making us wonder how fasting triggers AKT activation in *Fbp1^ΔHep^* mice. Postulating that the fasting induced increase in AKT activation, steatosis and hepatomegaly in *Fbp1^ΔHep^* mice could be due to FA-stimulated insulin secretion in response to fasting-induced lipolysis (Crespin et al., 1969), we injected *Fbp1^F/F^* and *Fbp1^ΔHep^* mice with high dose streptozotocin (STZ) to destroy insulin producing β cells (Graham et al., 2011). As expected, STZ treatment reduced blood insulin and increased blood glucose, which was slightly lower in *Fbp1^ΔHep^* mice than in *Fbp1^F/F^* mice after a 4h fast (Figures S7P and S7Q). Importantly, STZ treatment blocked fasting-induced hepatomegaly, hyperlipidemia, AKT and GSK3β phosphorylation, and ACLY, ACC1, FASN, HMGCS, HMGCR and CD36 mRNA expression (Figures S7R-S7Y), indicating that the liver phenotypes manifested by fasted *Fbp1^ΔHep^* mice depend on insulin secretion from β cells.

### A complex disrupting FBP1-derived peptide ameliorates insulin resistance

In addition to deletion and nonsense mutations that block FBP1 expression, human FBP1 deficiency can be caused by missense mutations (Emecen Sanli et al., 2022; Salih et al., 2020). We stably expressed FBP1 missense mutants in Huh7 cells and found most of them to be poorly expressed and devoid of catalytic activity (Figures 7A and S8A). Those variants that were expressed, R158W, G164S, N213K and L329P, no longer inhibited insulin stimulated AKT phosphorylation (Figure 7A). Particularly interesting is FBP1^L329P^ in exon 7, which was relatively well expressed with only a partial loss of FBP1 catalytic activity but completely devoid of AKT inhibitory activity (Figures 7A, S8A and S8B). Unlike WT FBP1 and FBP1^E98A^, FBP1^L329P^ did not associate with AKT, ALDOB and PP2A-C (Figure 7B). To further delineate FBP1 sequences mediating these interactions we used FBP1 deletion mutants, ΔE1-ΔE7 (Li et al., 2014), each lacking one *FBP1* exon (E) (Figure S8B). ΔE6 and ΔE7 did not bind AKT nor inhibited its activity and ΔE7 did not interact with PP2A-C, but ΔE1 was only defective in ALDOB binding (Figures 7C and S8C). These results suggest that FBP1 assembles the AKT inhibitory complex through separate and distinct contacts with AKT, PP2A-C and ALDOB. To test whether a synthetic E7 peptide that binds AKT or PP2A-C can disrupt the FBP1:PP2A-C:ALDOB:AKT complex and lead to AKT activation, we synthesized a peptide encompassing FBP1 AA 275 to 300 preceded by a TAT-derived cell penetrating peptide (Figure S8D). According to AlphaFold structural prediction and docking analysis, the E7 peptide is alpha helical, assuming the same fold it has within native FBP1, and is capable of differential association with the N-termini of AKT1 and PP2A-C but not with ALDOB (Figures S8E-S8G).

**Figure 7.**
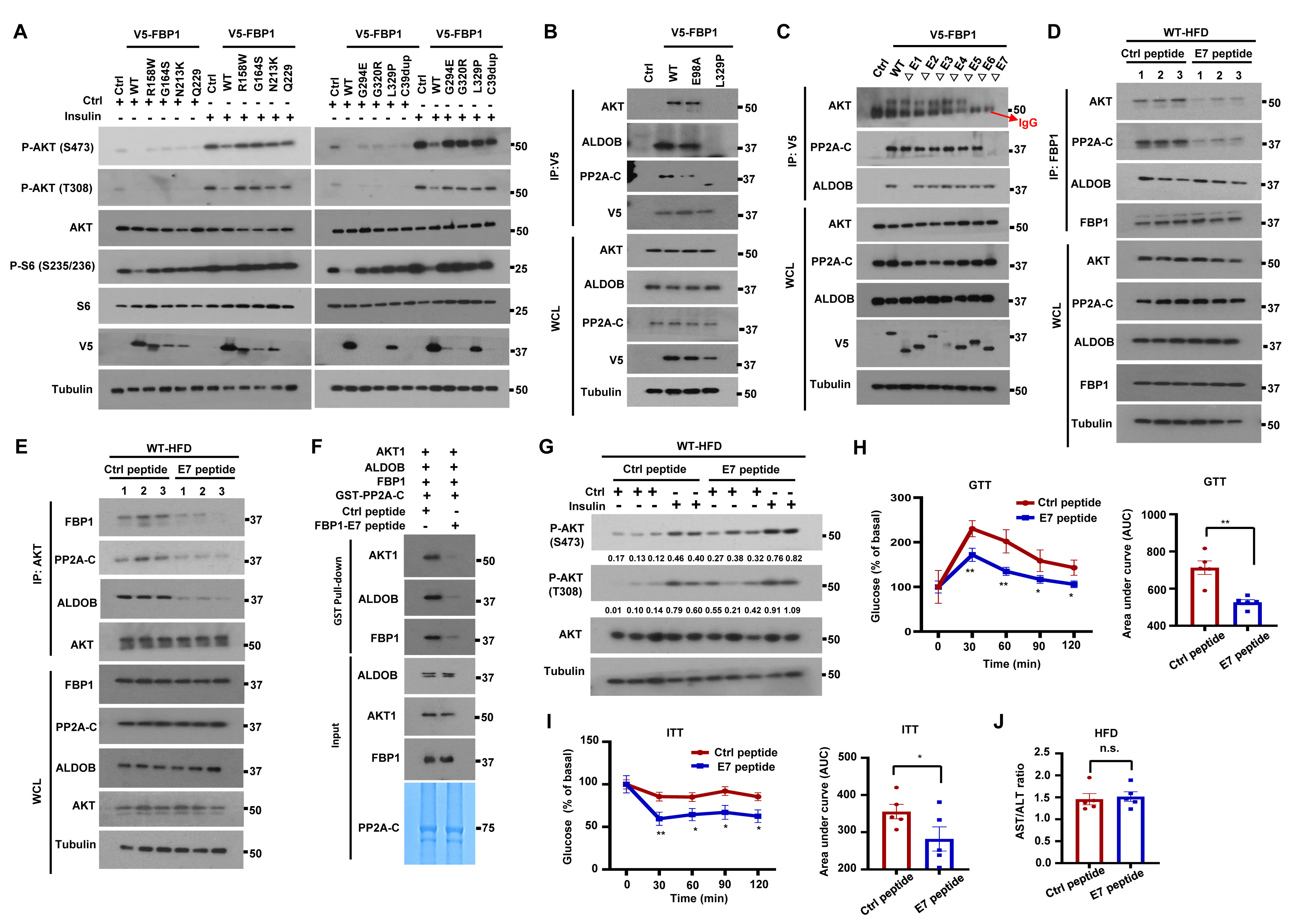
Complex-disrupting FBP1-derived C-terminal peptide ameliorates insulin resistance. (A) IB analysis of Huh7 cells stably expressing V5-tagged human FBP1 missense mutants incubated -/+ insulin (100 nM, 1 h) after 6 h serum starvation. (B) V5-FBP1, V5-FBP1^E98A^ and V5-FBP1^L329P^ were IP’ed from stably transfected Huh7 cells and the gel separated IPs were IB’ed with the indicated antibodies. (C) To map FBP1 regions interacting with AKT, PP2A-C and ALDOB, V5-tagged FBP1 exon deletion mutants were stably expressed in Huh7 cells and their association with AKT, PP2A-C and ALDOB was examined by IP with V5 antibody. (D and E) Male BL6 mice were placed on HFD for 14 w to induce insulin resistance. The mice were injected every other day with Ctrl or FBP1 E7 peptides (10 mg/kg each) for 2 w. The mice were then fasted overnight before analysis. Liver lysates were IP’ed with FBP1 (D) or AKT (E) antibodies and AKT, FBP1 and PP2A-C co-IPs were IB analyzed. (F) The GST-PP2A-C pull down described in Figure 6A was performed in the presence of FBP1 E7 or Ctrl peptides and analyzed as above. (G) HFD-fed BL6 mice from 7D were injected -/+ insulin 15 min before being analyzed for liver AKT activation. Relative normalized protein amounts are indicated below the P-AKT strips. (H) Mice from 7D were fasted overnight and subjected to GTT (n=5) (left) and AUC quantification (right). (I) ITT of mice from 7D that were fasted for 2-4 h (n=5) (I, left) and AUC quantification (I, right). (J) AST/ALT ratio in mice from 7D (n=5). Data are presented as mean ± SEM. *P < 0.05, **P < 0.01 (Unpaired two-tailed t test).

As *FBP1* mRNA is highly expressed in obese and T2D patients (GSE15653 and GSE64998) (Figures S8H-S8J), we examined whether FBP1:PP2A-C:ALDOB:AKT complex disruption reverses insulin resistance. We injected the E7 peptide or a scrambled peptide to HFD-fed BL6 mice. Strikingly, the C-terminal peptide disrupted the association of FBP1 with AKT and PP2A-C, and AKT with FBP1 and PP2A-C (Figures 7D and 7E). GST pulldown and PLA confirmed that the FBP1 E7 peptide reduced the association of PP2A-C with FBP1, AKT and ALDOB, as well as the AKT-PP2A-C, AKT-FBP1 and FBP1-PP2A-C interactions, but had no effect on the FBP1-ALDOB interaction (Figures 7F and S8K-S8N), which is mediated by FBP1 E1. The E7 peptide enhanced basal and insulin-stimulated AKT activation in HFD-fed WT mice (Figure 7G). In vitro dephosphorylation assays showed E7 peptide retarded AKT dephosphorylation by PP2A-C (Figure S8O). Importantly, E7 peptide treatment restored glucose tolerance and insulin sensitivity to HFD-fed mice, modestly reducing gluconeogenesis, lowering their blood glucose and increasing liver glycogen and lipids with no effect on serum TG, weight gain or FBP1 catalytic activity (Figures 7H-7J and S8P-S8W). These results establish FBP1 as an important attenuator of insulin signaling in WT mice, showing that this non-enzymatic FBP1 function can be exploited in the development of insulin sensitizers for treatment of obesity-related insulin resistance.

## DISCUSSION

Until now, the sole metabolic function assigned to FBP1 is the conversion of F1,6-P_2_ to F6-P, which controls GNG (Hunter et al., 2018; Leithner, 2020). In cancer cell nuclei, however, FBP1 was found to inhibit HIF1α activity through protein-protein interactions (Li et al., 2014). Here we show that cytoplasmic FBP1 has another critical and physiologically relevant metabolic function, the formation of a regulatory complex containing PP2A-C and ALDOB that binds AKT and blunts its activation. Discovery of this heretofore unrecognized function, in which FBP1 serves as a “safety valve” that prevents excessive insulin signaling and balances glucose and lipid metabolism, was enabled by generation of an accurate mouse model of human FBP1 deficiency (OMIM;229700). Like FBP1 deficient infants, *Fbp1^ΔHep^* mice, which lack hepatic FBP1 since an early age, are generally normal but exhibit rapid and severe hypoglycemia, hepatomegaly, hepatosteatosis, hyperlipidemia and liver damage upon fasting. Hypoglycemia and low liver glycogen stores are due to lack of FBP1 catalytic activity, which is needed for GNG and generation of G6-P, the allosteric activator of glycogen synthase. By contrast, the liver pathologies and hyperlipidemia depend on insulin stimulated AKT hyperactivation and not on FBP1 catalytic activity. Accordingly, these pathologies, and not hypoglycemia, are prevented by re-expression of catalytically inactive FBP1, AKT inhibition and to a lesser extent by inhibition of mTORC1, the AKT effector that stimulates protein synthesis (Bhaskar and Hay, 2007), a likely driver of hepatomegaly. mTORC1 also activates the lipogenic transcription factor SREBP1c (Bakan and Laplante, 2012), accounting for increased lipogenesis and hyperlipidemia. Enhanced AKT activation may also contribute to some of the long-term metabolic abnormalities seen in older FBP1 deficient individuals (Gorce et al., 2022). Of note, hepatic FBP1 ablation in older (24-wo) Fbp1^F/F^ mice using AAV-CRE transduction resulted in pronounced liver damage, fibrosis and enhanced hepatocyte tumorigenesis, with a negligible effect on glucose tolerance (Li et al., 2020). These differences are most likely due to the timing of FBP1 ablation and the evaluation of glucose tolerance in lean vs. HFD-fed mice.

While causing AKT hyperactivation in fasted and insulin-treated mice, hepatocyte FBP1 deficiency protects HFD-fed mice from glucose intolerance. The catalytically inactive FBP1^E98A^ variant binds ALDOB and PP2A-C and inhibits AKT activation as effectively as the native protein, but human *FBP1* missense mutants that are not severely destabilized, L329P, R158W, G164S and N213K, are devoid of AKT inhibitory activity. One of the residues affected by these mutations, L329, is not a part of the catalytic pocket (Huang et al., 2020; Ruf et al., 2016) and is associated with only a modest decrease in FBPase activity, underscoring the disconnect between loss of GNG and insulin hyperresponsiveness. Mass spectrometry, IP, gel filtration and PLA demonstrated that FBP1 is the lynchpin responsible for formation of the FBP1:AKT:PP2A-C:ALDOB regulatory complex that is destabilized by insulin to facilitate AKT activation. HFD stimulates FBP1 expression and FBP1:AKT:PP2A-C:ALDOB complex formation, suggesting that elevated FBP1 expression is a previously unknown contributor to HFD-induced insulin resistance. Although more precise understanding of the interactions between these proteins requires detailed structural analysis, we found that a synthetic peptide derived from *FBP1* E7 can be docked into AKT and PP2A-C and disrupt their interactions with native FBP1. By disrupting the AKT inhibitory complex this peptide serves as an AKT activator and an insulin mimic, capable of reversing obesity-induced glucose intolerance, further demonstrating the physiological relevance of the FBP1 nucleated AKT inhibitory complex. Although like insulin secretagogues and other anti-diabetic drugs E7 treatment modestly potentiates hepatosteatosis, the restoration of insulin sensitivity outweighs the increased risk of hepatosteatosis, which could be circumvented by co-treatment with β oxidation inducing uncouplers (Goedeke et al., 2019). Our biochemical analysis suggests that the metabolism regulating function of FBP1 depends on the recruitment of PP2A-C to AKT, rather than other phosphoproteins, thus attenuating AKT activation by insulin. Irreversible and complete loss of this function, which probably appeared early in vertebrate evolution, as it modulates liver energy storage and lipid production that are highly important in egg laying animals, results in dangerous insulin hyperresponsiveness, further compounded by the absence of GNG.

### Limitations of the study

While our study indicates that FBP1 inhibits insulin hyperresponsiveness by promoting assembly of the FBP1:AKT:PP2A-C:ALDOB regulatory complex, which can be assembled in vitro from purified proteins, we cannot rule out the presence of additional components of this complex. We have used FBP1 ablation to demonstrate the importance of its AKT regulating function in mice and human hepatocytes, but due to the absence of human clinical specimens we did not demonstrate that the human FBP1 deficiency also results in AKT hyperactivation. The E7 peptide reverses insulin resistance by disrupting the FBP1:AKT:PP2A-C:ALDOB complex, but further sequence optimization may create more potent E7 variants that will ameliorate insulin resistance at a lower dose. Clinical trials will be needed for establishing the human efficacy of such peptides and determine whether they can reverse insulin resistance without further aggravation of hepatosteatosis.

## STAR★METHODS

Detailed methods are provided in the online version of this paper and include the following:

● KEY RESOURCES TABLE
● RESOURCE AVAILABILITY

○ Lead contact
○ Materials availability
○ Data and code availability
● EXPERIMENTAL MODEL AND SUBJECT DETAILS

○ Mice
○ Cell Culture and Reagents
○ Human hepatocytes
● METHOD DETAILS

○ Constructs
○ Plasmid transfection and virus infection
○ Immunoprecipitation and immunoblot analysis
○ Histology
○ Measurements of metabolites and hormones
○ [U-^13^C] glucose, ^13^C-lactate and D2O labeling
○ Stable-Isotope Tracer Metabolomics
○ Size Exclusion Chromatography
○ FBP1 activity assay
○ GTT, PTT and ITT
○ Glycogen extraction
○ Glycogen synthase activity
○ Measuring liver protein synthesis with O-propargyl-puromycin
○ Generation and infection by AAV8-FBP1, and AAV8-FBP1^E98A^ and AAV8-Ctrl virus
○ Primary mouse hepatocytes isolation
○ Proximity ligation assays (PLA)
○ Mass spectrometry
○ In vitro binding assay
○ In vitro PP2A dephosphorylation assay
○ Peptide synthesis and treatment
○ Protein structure predictions and peptide docking
○ RNA isolation and quantitative real-time PCR (Q-PCR)
● QUANTIFICATION AND STATISTICAL ANALYSIS

## SUPPLEMENTAL INFORMATION

Supplemental information can be found with this article online.

## ACKNOWLEDGEMENTS

We thank the Karin Lab for helpful discussions and Cell Signaling Technologies, Santa Cruz Technologies, and Life Technologies for gifts of antibodies/other reagents, the UCSD histology core for assistance, Sanford Burnham Prebys Cancer Metabolism Core for the metabolomic analyses and Collabotarive Center for multiplexed proteomics at UCSD. Funding: Research was supported by NIH grants to M.K. (R01DK120714, R01CA234128), M.K. and J.H.L. (R01DK133448), M.C.S. (P01CA104838 and R35CA197602), A.R.S. (R01DK117551, R01DK125820, R01DK076906, and P30DK063491), W.Y. (R21HD107516, R00DK115998, and R01DK125560) and A.C.N. (R35 GM122523). M.K. hold the Ben and Wanda Hildyard Chair for Mitochondrial and Metabolic Diseases and is an American Cancer Society Research Professor. A. C. J. was supported in part by the University of California, San Diego, Graduate Training Program in Cellular and Molecular Pharmacology (T32 GM007752) and the National Science Foundation Graduate Research Fellowship (#DGE-1650112).

## AUTHOR CONTRIBUTIONS

M.K. and L.G. conceived the project. L.G. and Y.H.Z. designed the study and performed most of the experiments. K.W., M. L., J.L.L., S. P., M.T., F.M.L., K.C.R.and A.C.J assisted with experiments and data analysis. J.M. assisted with gel filtration analysis. B.C.Z. performed glycogen content anaylsis and GYS2 IB. L.C.K selected and provided the FBP1^E98A^ variant. X.L and T.K. provided human hepatocytes. J.H.L. performed Seq-Scope analysis. I.H.W. and D.J.G. performed the mass spectrometry. W.Y., A. C. N., A.R.S. and M.C.S. provided important reagents, advice and helpled with data interpretation and discussion. M.K. and L.G. wrote the manuscript, with all authors contributing and proving feedback and advice.

## DECLARATION OF INTERESTS

M.K. and A.R.S. are founders and stockholders in Elgia Pharmaceuticals and M.K. had received research support from Merck and Janssen Pharmaceuticals. All other authors declare no competing interests.

## STAR★METHODS

### KEY RESOURCES TABLE

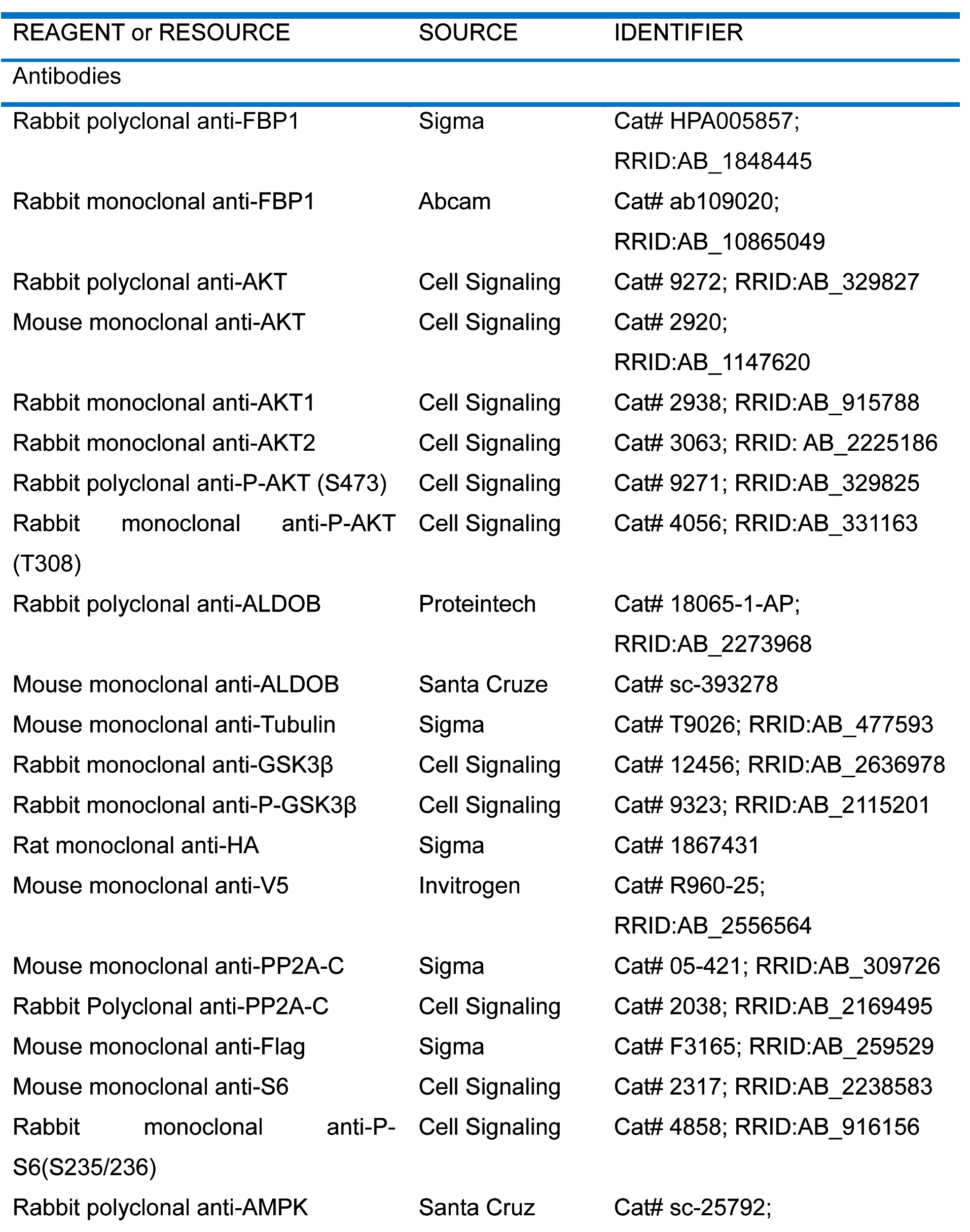
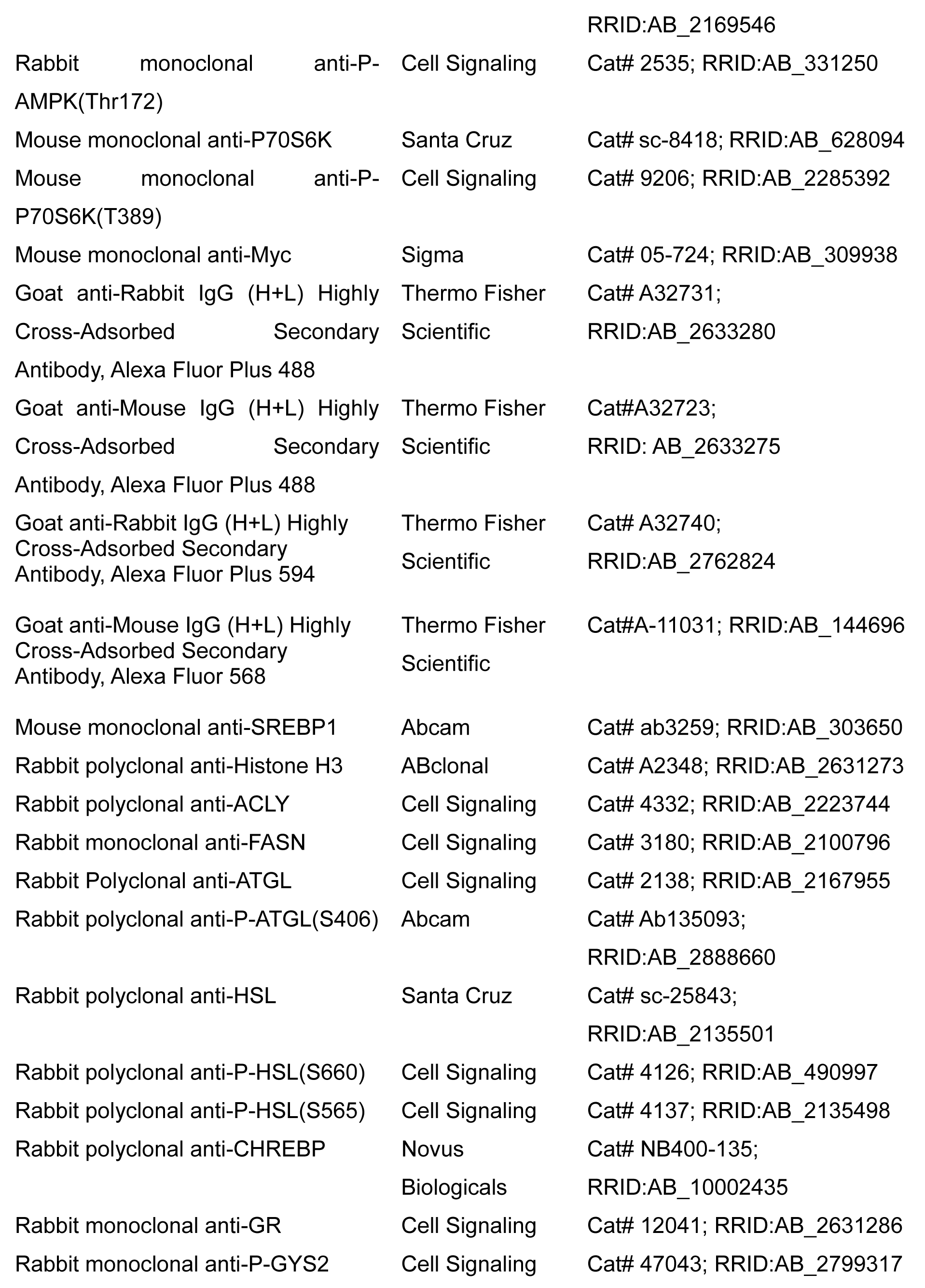
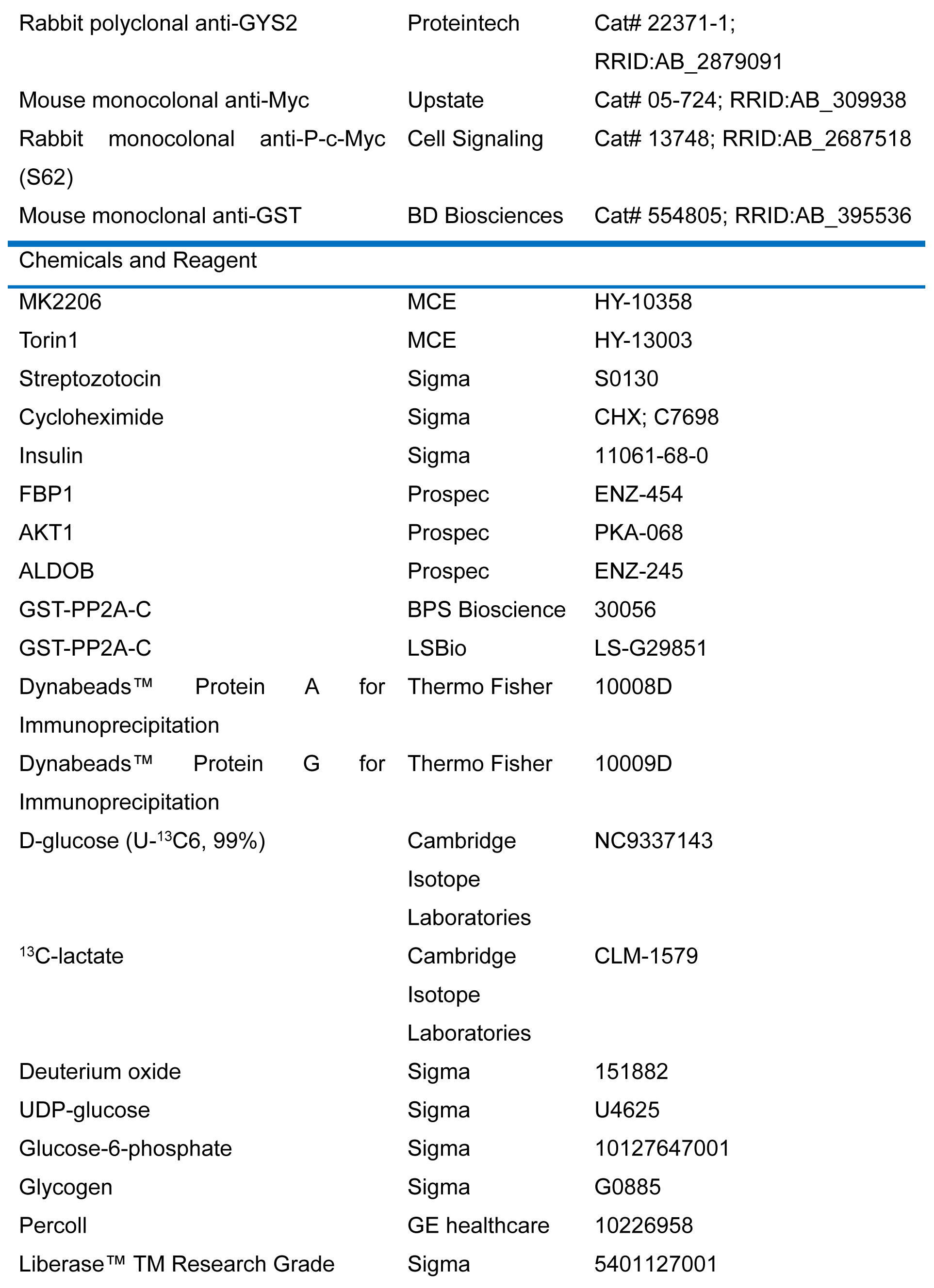
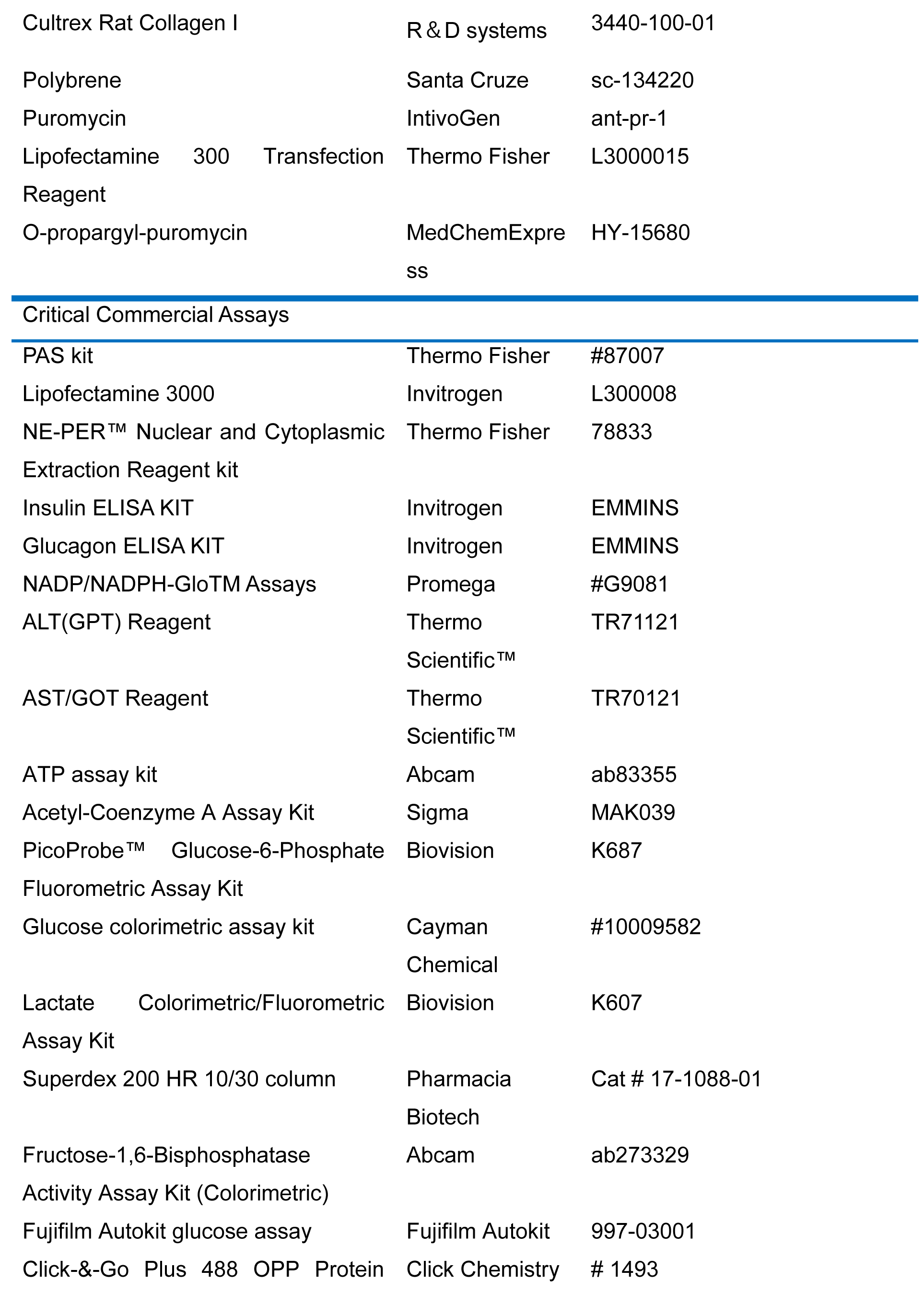
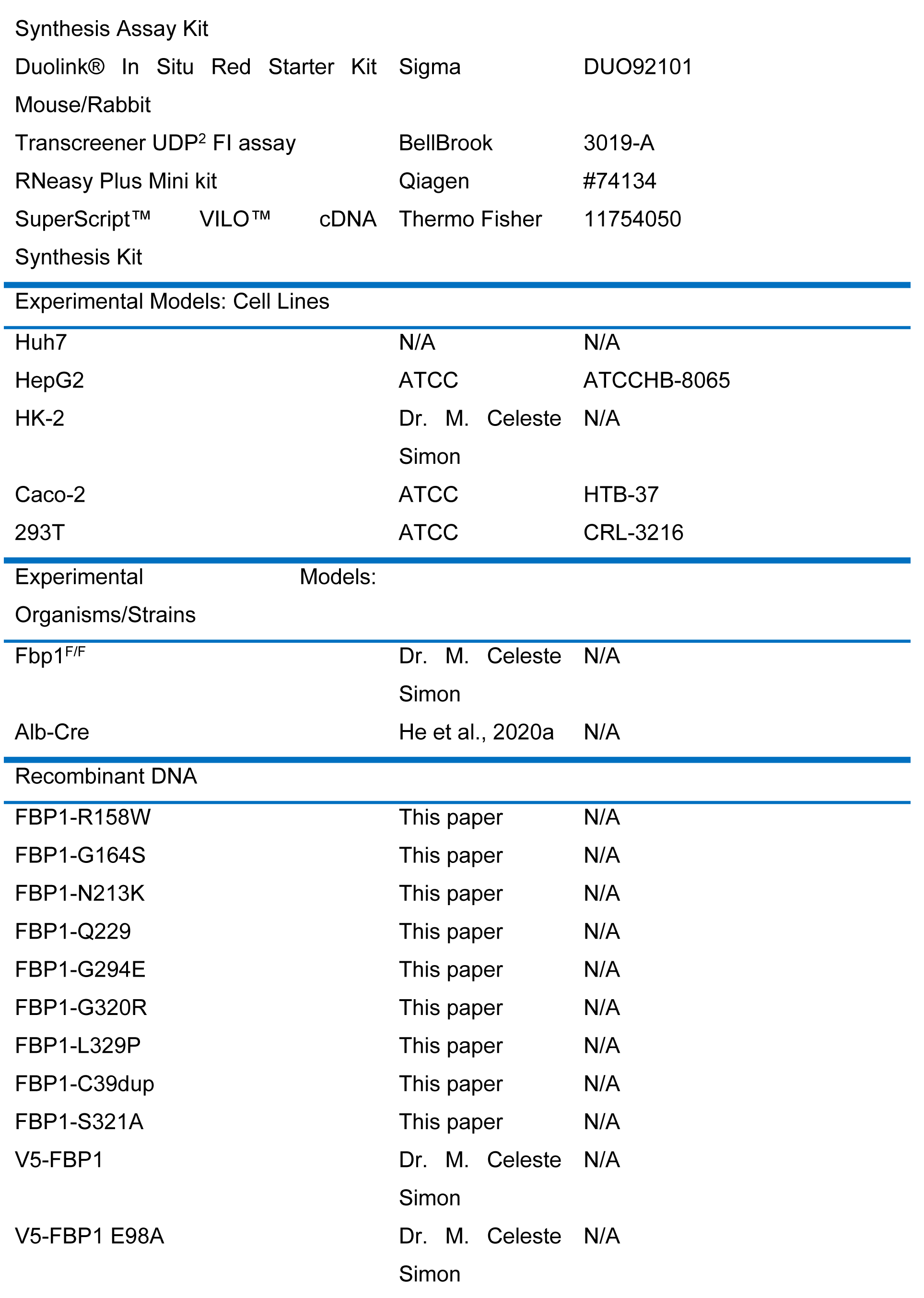
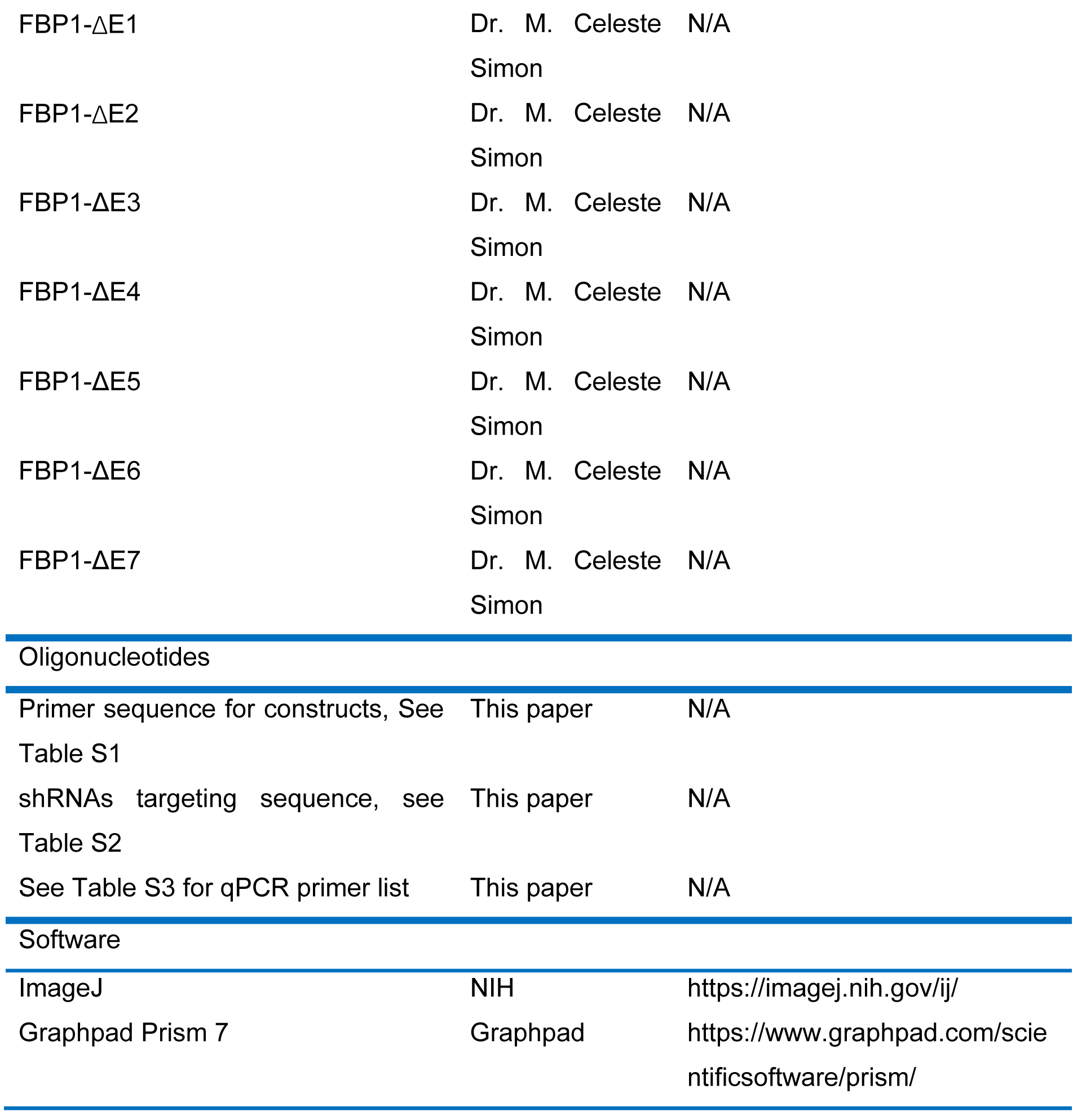

## RESOURCE AVAILABILITY

### Lead contact

Further information and requests for resources and reagents should be directed to and will be fulfilled by the Lead Contact, Michael Karin.

### Materials availability

This study generate new unique reagents is available from the lead contact upon request.

### Data and code availability

● All data reported in this paper will be shared by the lead contact upon request. Source data and uncropped western blots for the figures in the manuscript are provided in Data S1.
● This paper does not report original codes.
● Any additional information required to reanalyze the data reported in this paper is available from the lead contact upon request.

## EXPERIMENTAL MODEL AND SUBJECT DETAILS

### Mice

*Fbp1^F/F^* mice were generated by Dr. M. Celeste Simon (University of Pennsylvania, Philadelphia) (Li et al., 2020) and crossed to *Alb-Cre* mice to generate *Fbp1^ΔHep^* mice at UCSD. All mice were backcrossed into the BL6 background at least nine generations, and only male mice were used in most experiments. Mice were maintained in filter-topped cages on autoclaved food and water with a 12 h light (6am-6pm)/ dark (6pm-6am) cycle. Experiments were performed according to UCSD Institutional Animal Care and Use Committee and NIH guidelines and regulations. Dr. Karin’s Animal protocol S00218 was approved by the UCSD Institutional Animal Care and Use Committee. Where indicated, mice were fed with HFD (#S3282, Bio-serv) for a total of 12 wk, starting at 7 wk-of-age. Body weight and food consumption were monitored bi-weekly throughout the entire feeding period (Kim et al., 2018). Mice were starved for 4 h or 16 h before blood collection and sacrifice and liver and adipose tissue were excised and weighed. Where indicated, male *Fbp1^F/F^* and *Fbp1^ΔHep^* mice (8 wo) were intraperitoneally (i.p.) injected with MK2206 (100 mg/kg) or Torin1 (10 mg/kg) on 2 sequential days prior to fasting from 6 pm to 10 am, at which point the mice were euthanized for blood and tissue collection. The number of mice per experiment and their ages are indicated in the figure legends.

For streptozotocin treatment, 8-10 wo *Fbp1^F/F^* and *Fbp1^ΔHep^* male and female mice were i.p. injected with 150 mg/kg STZ. Mice were monitored daily for food and water consumption. Blood glucose was measured daily at random. After 5-7 days, mice with substantial hyperglycemia were fasted for 4 h before euthanasia and liver and blood analyses.

### Cell Culture and Reagents

Human HCC HepG2 and Huh7 cell lines were purchased from ATCC (HB-8065) and Cell bank, respectively. Huh7, HepG2, Caco-2 and 293T cells were tested regularly to confirm they are mycoplasma negative, and cultured in low glucose DMEM (Life technologies, 11885084), MEM (Life technologies, 11095080, and DMEM (Life technologies, 2366044), respectively, plus 10% fetal bovine serum (FBS) (Gibco), penicillin (100 mg/ml) and streptomycin (100 mg/ml). The HK-2 kidney cell line was provided by Dr. Celeste Simon and cultured in Keratinocyte-SFM medium supplemented with human recombinant epidermal growth factor (EGF) and bovine pituitary extract (BPE) (Life Technologies)(Li et al., 2014). Cells were incubated at 37℃ in a humidified chamber with 5% CO2. Cells were treated with insulin (Sigma 11061-68-0) at the times and concentrations indicated in figure legends.

### Human hepatocytes

Donor livers rejected for transplantation were obtained via Lifesharing OPO as a part of T. Kisseleva’s research program. Donors, who had no history of alcohol abuse, body-mass index 33.82, had no liver fibrosis and minimal liver steatosis, was qualified in this study conducted under IRB171883XX (approved on November 2017 by UCSD human Research Protection Program, under the title “Unused liver from deceased donors: role of myofibroblasts in liver fibrosis”). Hepatocytes isolation and purification were as described previously (Todoric et al., 2020). The hepatocytes were plated in collagen coated 6-wells and after attached, were incubated with shCtrl and shFBP1 lentivirus for 24 h. After recovery for 24-48 h, the cells were stimulated with insulin for the indicated times and concentrations after 6 h of serum starvation.

## METHOD DETAILS

### Constructs

V5-FBP1, V5-FBP1^E98A^ and FBP1 truncation mutations (ΔE1-ΔE7) were generated in Dr. M. Celeste Simon’s lab. Flag-ALDOB, Flag-PP2A-C, HA-AKT1 were kindly provided by Drs. Huiyong Yin (Chinese Academy of Sciences, Shanghai, China) and Alexandra C. Newton (University of California at San Diego, USA), respectively. FBP1 (TRCN0000050034, TRCN0000050035) and PP2A-C (TRCN0000002483, TRCN0000002486) shRNAs were purchased from Sigma. SgALDOB was constructed by cloning the guide sequences into the BsmBI site of the lentiCRISPR v2-puro vector. Human FBP1 mutants were constructed by Q5 Site-Directed Mutagenesis Kit (NEB, E0554S) in the PCDH-V5-FBP1 vectors. Primers are listed in Tables S1 and S2.

### Plasmid transfection and virus infection

Expression vectors were transfected into 293T cells or HCC cell lines using Lipofectamine 3000 (Invitrogen, L300008) following the manufacturer’s protocol. Lentiviruses were produced by co-transfecting PSPAX2 (4 μg), PMD2.G (4 μg) and PCDH-V5-FBP1 (8 μg) or FBP1 mutants (8 μg), PLKO.1 (8 μg) or lentiCRISPERv2 (8 μg) vectors. After 8 h medium was changed and fresh DMEM plus 10% FBS was added. Virus containing media were harvested 48-64 h later and filtered by 0.45 μm Steriflip filter (Millipore). HepG2 and Huh7 cells were infected with 1 ml virus containing medium with 8 μg/ml polybrene for 24 h. Cells were allowed to recover in complete medium for 24-48 h and then selected with puromycin. After 48 h surviving cell pools were used in the different experiments.

### Immunoprecipitation and immunoblot analysis

Cells were harvested and lysed in RIPA buffer (50 mM Tris-HCl, pH 7.4, 150 mM NaCl, 1% Triton X-100, 1% sodium deoxycholate, 0.1% SDS, 1 mM EDTA) supplemented with complete protease inhibitor cocktail (Gu et al., 2021). Livers were homogenized in a Dounce homogenizer (Thomas Scientific, NJ) with 30 strokes in RIPA buffer with complete protease inhibitor cocktail. For immunoprecipitation (IP) experiments, cells were harvested in lysis buffer (20 mM Tris-HCl, pH 7.5, 1% NP-40, 137 mM NaCl, 1 mM MgCl_2_, 1 mM CaCl_2_, 10% glycerol) supplemented with complete protease inhibitor cocktail. Cell lysates were pre-cleared with 30 μl protein G beads (Life Technologies), and then incubated with 2 μg isotype matched IgG control or indicated antibodies on a rocking platform overnight at 4℃. 50 μl protein G were added and incubated for another 2-3 h. The immunocomplexes were washed 5x with lysis buffer, separated by SDS-PAGE and transferred to polyvinylidene difluoride (PVDF) membranes, blocked in 5% nonfat milk, and incubated with the indicated primary antibodies overnight. Second antibodies were added for another 1 h and detected with Clarity Western ECL Substrate (Biorad). Immunoreactive bands were exposed in an automatic X-ray film processor. Nuclear extraction was performed with the NE-PER™ Nuclear and Cytoplasmic Extraction Reagent kit (Thermo Fisher, 78833), following manufacturer’s instructions. After extraction, nuclear and cytoplasmic extracts were separated by SDS-PAGE and analyzed by immunoblotting as above.

### Histology

Livers were dissected, fixed in 4% paraformaldehyde and embedded in paraffin. 5 μm thick sections were stained with hematoxylin and eosin (H&E) (Leica, 3801615, 3801571). For frozen block preparation, livers were embedded in Tissue-Tek OCT compound (Sakura Finetek), sectioned, and stained with Oil Red O to visualize TG accumulation. Liver PAS staining was performed with PAS kit (Thermo Fisher Scientific, #87007), using the manufacture’s protocol.

### Measurements of metabolites and hormones

Liver TG, serum TG and serum cholesterol were measured with Triglyceride Colorimetric Assay Kit (Cayman Chemical #10010303) and Cholesterol Fluorometric Assay Kit (Cayman Chemical #10007640), respectively, according to manufacturer protocols. Serum insulin and glucagon concentrations were determined by Mouse Insulin ELISA KIT (Invitrogen, EMMINS) and Glucagon Quantikine ELISA Kit (R&D Systems, DGCG0), respectively, following manufacturer protocols. Liver NADPH and NADP were measured by NADP/NADPH-Glo^TM^ Assays (Promega #G9081) according to manufacturer’s protocol. ALT and AST assays were performed with ALT(GPT) Reagent (Thermo Scientific™, TR71121) and AST/GOT Reagent (Thermo Scientific™, TR70121), respectively, according to the manufacturer’s protocol. ATP, Acetyl-CoA and G6P concentrations were determined by ATP assay kit (Abcam, ab83355), Acetyl-Coenzyme A Assay Kit (Sigma, MAK039) and PicoProbe™ Glucose-6-Phosphate Fluorometric Assay Kit (Biovision, K687), respectively according to the manufacturers protocols. Glucose and lactate were measured with Glucose colorimetric assay kit (Cayman Chemical #10009582) and Lactate Colorimetric/Fluorometric Assay Kit (Biovision, K607), respectively, according to the manufacturer’s protocols.

### [U-^13^C] glucose, ^13^C-lactate and D2O labeling

D-glucose (U-^13^C6, 99%) and ^13^C-lactate were purchased from Cambridge Isotope Laboratories (NC9337143, CLM-1579). 8 wo *Fbp1^F/F^* and *Fbp1^ΔHep^* mice were fasted overnight from 6 pm-10 am and 45 min before sacking [U-^13^C] glucose was treated as previously described (Sengupta et al., 2020; Zwingmann and Bilodeau, 2006). For lactate labeling, mice were treated with sodium L-lactate (^13^C3) 60 min before sacking as previously described (Gray et al., 2015). Liver and serum were collected for metabolomic analysis. To measure DNL, 8 wo *Fbp1^F/F^* and *Fbp1^ΔHep^* mice were i.p. injected with 0.035 ml/body weight D_2_O (Sigma) in 0.9% NaCl, and drinking water was replaced with 8% enriched D_2_O (Huh et al., 2020). Mice were given D_2_O for 22-24h and starved for 16 h before sacrifice, plasma and liver were collected and immediately snap-frozen in liquid nitrogen.

### Stable-Isotope Tracer Metabolomics

Sample extraction methods: Frozen liver samples of *Fbp1^F/F^* and *Fbp1^ΔHep^* (∼40 mg) were transferred to 2-mL tubes containing 2.8 mm ceramic beads (Omni International) and 0.45 ml ice-cold 50% methanol/ 20 µM L-norvaline was added. Tubes were shaken (setting 5.5) for 30 s on a Bead Ruptor 12 (Omni International), quickly placed on ice, and frozen at −80°C overnight. Thawed samples were centrifuged at 15,000 x g for 10 minutes at 4°C. The supernatant was then transferred to a new tube, mixed with 0.225 ml chloroform, and centrifuged at 10,000 x g for 10 minutes at 4°C. This produced a two-phase separation. Portions (100 µl) of the top phase were dried (Speedvac) for analyses of polar metabolites. The lower phase was dried (Speedvac) for analysis of fatty acids. Plasma samples (5 or 10 µl) were mixed with 4 volumes of acetone (pre-cooled to −20°C), left at −20°C for 1 h, and centrifuged at 14,000 x g for 10 minutes at 4°C. Supernatants were dried by Speedvac.

Metabolite derivatization and GC-MS run conditions: Polar metabolites except for glucose and sugar-phosphates were derivatized using isobutylhydroxylamine and MTBSTFA, and analyzed by GC-MS as described (Scott, 2021). Plasma samples for glucose analysis were derivatized first with 30 µl ethylhydroxylamine (Sigma) at 20 mg/ml in pyridine for 20 min at 80°C, and secondarily with 30 µl BSTFA (Thermo) for 60 min at 80°C. Samples were transferred to autosampler vials with inserts and analyzed using an Rxi-5ms column (15 m x 0.25 i.d. x 0.25 μm, Restek) installed in a Shimadzu QP-2010 Plus gas chromatograph-mass spectrometer (GC-MS). The GC-MS was programmed with an injection temperature of 250°C, 1.0 µl injection volume and split ratio 1/10. The GC oven temperature was initially 130°C for 4 min, rising to 230°C at 6°C/min, and to 280°C at 60°C/min with a final hold at this temperature for 2 min. GC flow rate, with helium as the carrier gas, was 50 cm/s. The GC-MS interface temperature was 300°C and (electron impact) ion source temperature was 200°C, with 70 eV ionization voltage. The main glucose peak eluted at 13.7 min, and fragments of m/z 319 (contains 4 glucose carbons; C_13_H_31_O_3_Si_3_) and m/z 205 (contains 2 glucose carbons; overall formula C_8_H_21_O2Si_2_) were used to analyze ^13^C-glucose labeling as described (Scott, 2021). Liver samples labeled with ^13^C-lactate for sugar-phosphate analysis were derivatized first with 30 µl pentafluorobenzyl-hydroxylamine (Alfa Aesar) 20 mg/ml in pyridine for 60 min at 37°C, and secondarily with 30 µl BSTFA (Thermo) for 60 min at 37°C. The GC-MS method was as described above, except that injection volume was 2 µl, and initial GC oven temperature was 200°C for 4 min, rising to 280°C at 8°C/min, with a final hold at this temperature for 2 min. F6-P eluted as a pair of peaks at 10.33 and 10.49 min. The earlier peak was analyzed for ^13^C-labeling (fragment of m/z 459 containing 3 glucose carbons; C_15_H_40_O_6_Si_4_P). Fatty acids (palmitate and stearate) were derivatized and analyzed for labeling as described (the methyl stearate ion m/z 298 was analyzed) (Scott, 2021).

### Size Exclusion Chromatography

Liver samples were fractionated on a Superdex 200 HR 10/30 column (Pharmacia Biotech, Cat # 17-1088-01) using an AKTA FPLC system (GE). Column was equilibrated with and samples were fractionated in CHAPS lysis buffer (25mM HEPES pH 7.4, 150mM NaCl, 1mM EDTA, and 0.3% CHAPS). 0.5 mL of sample at 10 mg/mL (5mg protein total) in CHAPS lysis buffer was injected onto the column and fractionated at a 0.5 mL/min flow rate over 1.2 column volumes. 0.25 mL fractions were collected. The injection loop was thoroughly rinsed, and blank runs were performed between each sample.

Molecular weights ranges within fractions were approximated through comparison with a standard curve (Kav of protein standards v. log(MW)). Kav of Gel Filtration Protein Standards (Bio-Rad, Cat#1511901) of known molecular weight were determined under identical fractionation conditions as described for samples and calculated using the following equation where Ve is elution volume, Vo is void volume, and Vt is the total volume of the column: Elution volumes were determined by monitoring inline absorbance at 280 nm as the standards eluted. Peak elution volumes were determined using Unicorn 5.1 Software (GE). Void volume was similarly determined by monitoring elution of 1 mg of blue dextran (Sigma D4772). Total volume of the column was provided by the manufacturer.

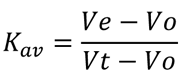

### FBP1 activity assay

FBP1 activity was measured with Fructose-1,6-Bisphosphatase Activity Assay Kit (Colorimetric) (Abcam, ab273329). Briefly, 20 μg lysates of control Huh7 cells, or Huh7 cells expressing WT and mutant forms of FBP1 were prepared in 500 μl ice-cold FBP assay buffer. (NH_4_)_2_SO_4_ was used to precipitate the proteins and remove small molecules that could interfere with the assay. The precipitated proteins were spun down and resuspend in FBP assay buffer. The samples were incubated with the reaction mix containing FBP converter, FBP probe, FBP developer and FBP substrate and absorbance at OD=450 was measured in a kinetic mode for 5-60 min at 37℃. FBP1 activity was normalized according to its expression level determined by immunoblotting.

### GTT, PTT, and ITT

For GTT and PTT, *Fbp1^F/F^* and *Fbp1^ΔHep^* mice that were HFD fed for 12 wk or NCD-8 wo mice were fasted for 12-14 h and then given 1 g/kg glucose or sodium pyruvate (2 g/kg, Sigma) by i.p. injection. Blood glucose was measured before injection and every 30 min thereafter with a glucometer (OneTouch Ultra 2, One Touch) on blood from superficial tail incision. 8 wo *Fbp1^F/F^* and *Fbp1^ΔHep^* mice or 18 wo *Fbp1^F/F^* and *Fbp1^ΔHep^* mice that were HFD fed for 12 wk, fasted for 2-4 h then injected with 0.5 U/kg insulin. Blood glucose was measured before injection and every 30 min thereafter, up to 2 h.

### Glycogen extraction

Small pieces of livers were weighted at dissection. Samples were boiled for 30 min in 500 μl of 30% KOH solution with Vortex every 10 min. 100 μl of 1 M Na_2_SO_4_ were added when the samples were cooled down. 1.2 ml of 100% ethanol was added, and the samples cooked for 5 min at 95℃. The samples were then centrifuged for 5 min at 16,000xg. Pellets were washed twice by resuspension in 500 μl of double-distilled water, 1 ml 100% ethanol, and centrifugation. Following the last wash, pellets were air-dried and resuspended in 100 μl of 50 mM sodium acetate at pH 4.8 containing 0.3 mg/ml amyloglucosidase. The samples were incubated at 37℃ overnight on a shaker to facilitate digestion. Fujifilm Autokit glucose assay (997-03001) was used to determine the amount of glycogen by comparing it to a standard curve. The content of glycogen was normalized to tissue weight.

### Glycogen Synthase Activity

Freeze-dried liver tissue (20g-30mg) was homogenized in ice-cold buffer (50 mM Tris-HCl (pH 7.8), 100 mM NaF, 10 mM EDTA). Homogenates were centrifuged for 5 min at 3600g at 4℃ and GYS activity measured in supernatants with 0 or 12 mM of added G6-P (Bouskila et al., 2010). Next, the Transcreener^®^UDP^2^ FI Assay Kit (BellBrook, 3019-A) was used to measure GYS activity by measuring the release of UDP from UDP-glucose.

### Measuring liver protein synthesis with O-propargyl-puromycin

8-wo *Fbp1^F/F^* and *Fbp1^ΔHep^* mice were starved for 4-6 h starting at 6 pm and then i.p. injected with 100 μl of a 20 mM solution of O-propargyl-puromycin (OP-Puro, MedChemExpress, HY-15680) in PBS (Liu et al., 2012). Livers were harvested after 1 h, fixed in formalin and paraffin embedded samples were cut into 5 μm sections. OPP incorporation was detected by staining with Click-&-Go Plus 488 OPP Protein Synthesis Assay Kit (Click Chemistry, # 1493), following manufacturer’s instructions. Briefly, liver sections were deparaffinized with xylene, hydrated with ethanol and incubated with a reaction cocktail containing copper catalyst, AZDye Azide plus solution and reducing agent for 20 min, while protected from light. DNA was stained with Hoechst 33342 and the sections imaged on a confocal microscope. The intensity of fluorescent-labeled protein-incorporated OPP was quantified by automated analysis of microscopic images with Image J software (NIH).

### Generation and infection by AAV8-FBP1, AAV8-FBP1^E98A^ and AAV8-Ctrl virus

pAAV[Exp]-CAG>mFbp1WT:T2A:EGFP:WPRE and pAAV[Exp]CAG>HA/mFbp1^E98A^:T2A: EGFP:WPRE were constructed by Vectorbuilder. AAV8-mFBP1, AAV8-mFBP1^E98A^ and AAV8-Ctrl viruses were generated by Vectorbuilder. Briefly, 293T cells were co-transfected with indicated vectors and Rep-cap plasmid and helper plasmids. After 50-60 h, cell lysates were harvested and concentrated by PEG precipitation and CsCl gradient ultracentrifugation. 6-8 wo mice were infected with AAV8-mFBP1, mFBP1^E98A^ and AAV8-Ctrl (10^12^ virus copies per mouse) via tail vein injection. Liver and serum were collected 3 weeks later.

### Primary mouse hepatocytes isolation

8-wo male *Fbp1^F/F^* and *Fbp1^ΔHep^* mice were fasted for 14-16 h and primary hepatocytes were isolated using a two-step collagenase perfusion as described (He et al., 2020a). Briefly, mice livers were perfused with perfusion buffer (HBSS (Thermo Fisher, 14175095) with 0.5mM EDTA and 25mM HEPES) and digested with digestion buffer (HBSS with Ca^2+^ Mg^2+^ (Thermo Fisher, 24020117), 25mM HEPES and 1mg/ml Liberase). The hepatocytes were spun down at 50g for 3 min at 4 ℃ then purified on a Percoll gradient. The hepatocytes were counted and directly collected or plated on collagen-coated plates for 6 h in plating medium (DMEM low glucose, 5%FBS and 1% Penicillin-streptomycin (PS)). After cell attachment, the plating medium was replaced with maintenance medium (Willams E medium, 2 mM glutamine and 1% PS) for further use.

### Proximity ligation assays (PLA)

AKT:FBP1:ALDOB:PP2A-C interactions in liver tissue were detected with an *in situ* PLA kit (Duolink® In Situ Red Starter Kit Mouse/Rabbit, Sigma, DUO92101) (Su et al., 2021). Briefly, paraffin embedded tissue sections were deparaffinized with 3x washes with xylene, rehydrated with ethanol and subjected to antigen unmasking with citrate buffer at 95-98 ℃ for 15 min. Tissues were incubated with 3% H_2_O_2_ for 10 min and then blocked with Duolink blocking solution at 37 ℃ for 60 min. The samples were incubated with primary antibodies overnight, and then incubated with the PLA Probe, ligase and polymerase following manufacturer’s instructions. Tissues were mounted with an *in situ* mounting medium with DAPI and images captured on a TCS SPE Leica confocal microscope. The results were quantified by counting dots per field in 3 randomly selected fields of view.

### Mass spectrometry

Sample Preparation: *Fbp1^ΔHep^* mouse livers transduced with AAV-FBP1 or AAV-Control were homogenized in NP-40 lysis buffer. Lysates were IP’ed with HA antibody using magnetic beads. The IP’s were washed twice with 1xPBS (Gibco) and resuspended in 150 uL 5% formic acid to denature proteins and remove them from the beads. The supernatants were placed in new tubes and 1.5 mL HPLC-grade water was added and then dried down in a concentrator (Thermo SpeedVac). Dried samples were resuspended in 250 uL 1M urea, reduced with 5 uL of 500 mM dithiothreitol (DTT) and placed on a 47℃ block for 30 minutes. Samples were then cooled to room temperature and 15 μL 500 mM iodoacetamide to alkylate proteins were added and the samples were kept in the dark for 45 minutes. The reactions were quenched with 5 uL 500 mM DTT. 1M Tris-HCl was added to adjust the pH and 5 μL trypsin (2.5 ug, Promega Sequence Grade) was added and the samples digested overnight at 37℃. 80 μL of 1% TFA was added to stop the digestion and the samples were dried down. The samples were then resuspended in 100 μL 0.1% formic acid (FA) desalted using C18 discs in a pipette tip (“Stage Tips”). The eluted peptides were then quantified (Thermo PepQuant) and 9 μg of peptides were transferred to a new tube and dried down. These samples were then resuspended in 9 μL 5% acetonitrile and 5% FA and were placed in MS inserts for proteomic characterization.

Mass spectrometry: Peptide samples were diluted to 1 μg/μl. Subsequently, 1 μl was loaded onto an in-house laser-pulled 100-μm-inner-diameter nanospray column packed to ∼300 mm with 1.8-μm 2Å C18 beads with a ∼1mm 5 μm frit (Reprosil). Peptides were separated via reverse-phase chromatography using a Thermo Easy-nLC 1000 liquid chromatography instrument (HPLC). Buffer A of the mobile phase contained 0.1% FA in HPLC-grade water, while buffer B contained 0.1% FA in ACN. An initial 2-min isocratic gradient flowing 3% B was followed by a linear increase up to 25% B for 85 min, increased to 45% B over 15 min, and a final increase to 95% B over 15 min, whereupon B was held for 6 min and returned to baseline (2 min) and held for 10 min, for a total of 183 min. The HPLC flow rate was 0.400 μl/min. Sample spectra were collected on a Thermo Fusion mass spectrometer that collected MS data in positive ion mode within the 400 to 1,500 m/z range.

Proteome Search: Mass spectra raw files were first searched using Proteome Discoverer 2.2 using the built-in SEQUEST search algorithm. Built-in TMT batch correction was enabled for all samples. Three FASTA databases were employed: Uniprot Swiss-Prot *Mus musculus* (taxon ID 10090). Target-decoy searching at both the peptide and protein level was employed with a strict FDR cutoff of 0.05 using the Percolator algorithm built into Proteome Discoverer 2.2. Enzyme specificity was set to full tryptic with static peptide modifications set to carbamidomethylation (+57.0214 Da). Dynamic modifications were set to oxidation (+15.995 Da) and N-terminal protein acetylation (+42.011 Da). Only high-confidence proteins (q < 0.01) were used for analysis.

### In vitro binding assay

In vitro binding assay was performed as described previously (He et al., 2020b). In brief, recombinant GST-PP2A-C protein were incubated with His-AKT, His-ALDOB and His-FBP1 in binding buffer (25mM Tris-HCl, 200 Mm NaCl, 1mM EDTA, 0.5% NP-40, 10μg/μl BSA and 1mM DTT). GST, AKT or FBP1 antibodies and protein A/G beads were mixed and incubated overnight. The lysates were washed with ice-cold binding buffer for 3 times and boiled before subjected to IB analysis.

### PP2A dephosphorylation assay

The PP2A dephosphorylation assay were performed as described (He et al., 2020b). Briefly, Huh7 cells stably expressing HA-AKT1 or HA-c-Myc were serum starved for 6 h and stimulated with insulin (100nM) or EGF (100ng/ml) for 1 h, respectively. Cells were harvested and AKT1 or c-Myc was IP’ed with an HA antibody. The immunoprecipitates were washed and resuspended in PP2A phosphatase assay buffer (20mM HEPES, 100mM NaCl and 3mM DTT). Together with recombinant ALDOB, FBP1 or FBP1^E98A^ and active PP2A, the AKT1or c-Myc IPs were incubated at 30 ℃ for the indicated times and AKT1 and c-Myc phosphorylation was assessed by IB analysis.

### Peptide synthesis and treatment

Cell permeable Scrambled (Ctrl) and FBP1 peptides were synthesized by Biomatik. Male BL6 mice were HFD fed for 14 weeks from week 7 postnatally. Mice were i.p. injected with the Ctrl and FBP1 peptides (10mg/kg each) every 2 days for 2 weeks. Mice were fasted overnight before being subjected to GTT and PTT and fasted 2h before ITT.

### Protein structure predictions and peptide docking

AKT1, PP2A-C, ALDOB, FBP1 and FBP1 E7 peptide structure modeling were performed with the ColabFold implantation of AlphaFold (https://alphafold.ebi.ac.uk/) (Jumper et al., 2021). The best positions for peptide and protein interactions were explored, and hydrogen bonds and amino acids were identified and labeled in PyMOL (https://pymol.org/2/).

### RNA isolation and quantitative real-time PCR (Q-PCR)

Total liver RNA was extracted with RNeasy Plus Mini kit (Qiagen #74134) and cDNA was synthesized with SuperScript™ VILO™ cDNA Synthesis Kit (Thermo Fisher Scientific, 11754050) (He et al., 2020a). mRNA expression was determined by CFX96 thermal cycler (Biorad). Data are presented as arbitrary units and calculated by comparative CT method (2Ct^(18s rRNA–gene of interest)^). Primers are listed in Table S3.

### QUANTIFICATION AND STATISTICAL ANALYSIS

Data are presented as mean ± SD or mean ± SEM as indicated. Differences between mean values were analyzed by unpaired two-tailed Student’s t test with GraphPad Prism software. P value<0.05 was considered as significant (*p < 0.05, **p < 0.01, ***p < 0.001, ****p < 0.0001, n.s., p≥0.05).

