## supplementary data for "FBP1 is a nonenzymatic safety valve that curtails AKT activation to prevent insulin hyperresponsiveness"

Li Gu *et al.*

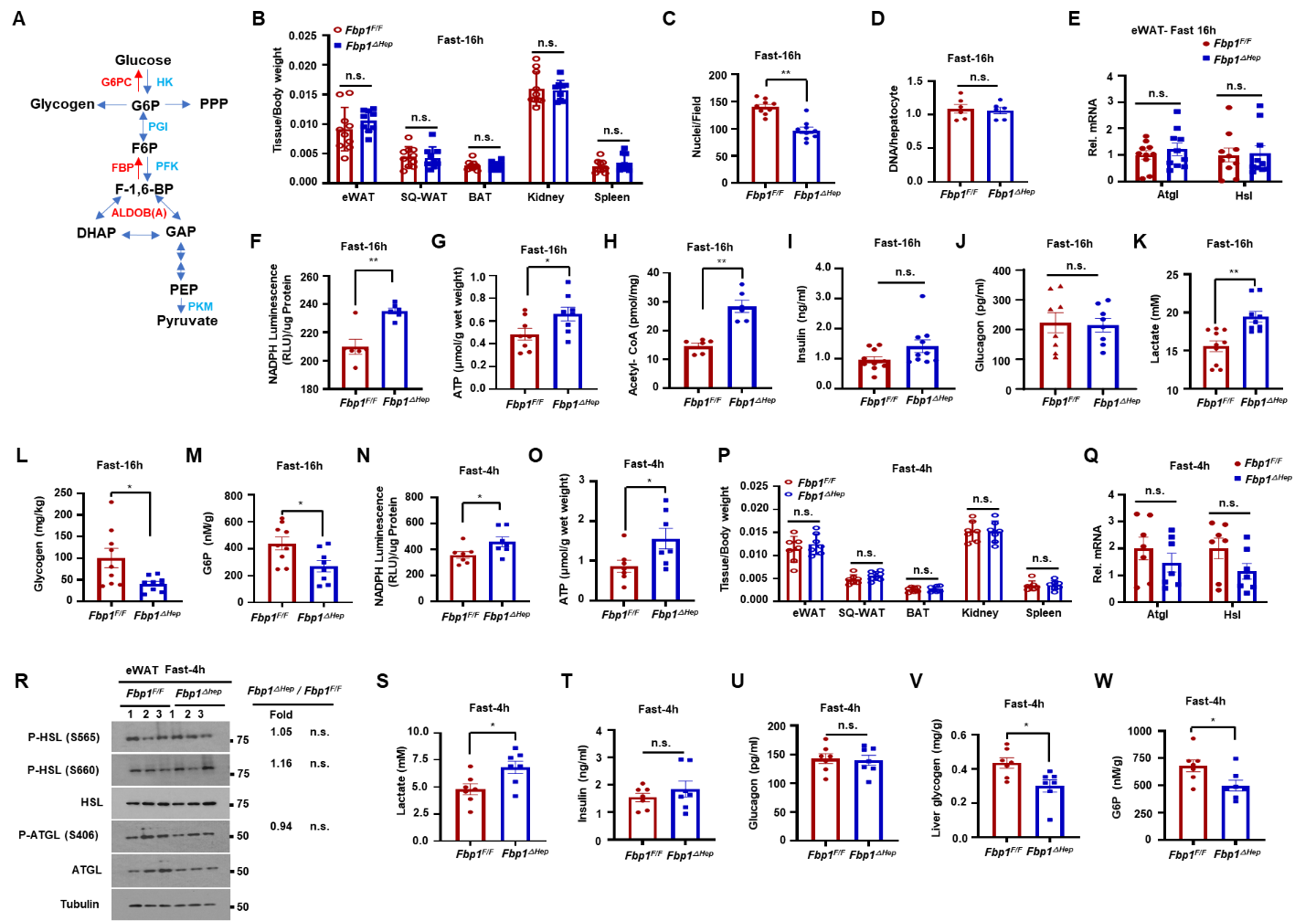

**Figure S1. Hepatocyte specific FBP1 ablation increases ATP, acetyl-CoA and serum lactate while decreasing glycogen and G6-P amounts after 16h and 4h fast, related to Figure 1.**

(A) Schematic representation of glucose metabolism and roles of FBP1 and ALDOB in GNG.

(B) Epididymal white adipose tissue (eWAT), subcutaneous adipose tissue (SQ-WAT), brown adipose tissue (BAT), kidney and spleen weight/body weight ratio in 16 h fasted 8-wo *Fbp1^F/F^* and *Fbp1^ΔHep^* mice (n=9-10).

(C and D) Nuclei per field (C) and hepatocyte DNA content (D) from 16 h fasted 8-wo *Fbp1^F/F^* and *Fbp1^ΔHep^* livers (n=6-9).

(E) qRT-PCR analysis of lipolytic mRNAs in eWAT of 16 h fasted 8-wo *Fbp1^F/F^* and *Fbp1^ΔHep^* mice (n=9-10).

(F-H) NADPH (F), ATP (G) and acetyl-CoA (H) concentrations of liver lysates from above mice (n=6-8).

(I and J) Serum insulin (I) and glucagon (J) in indicated mice (n=6-8).

(K-M) Serum lactate (K), liver glycogen (L) and G6-P (M) in indicated mice (n=9-10).

(N and O) NADPH (N) and ATP (O) concentrations in liver lysates of 4 h fasted 8-wo *Fbp1^F/F^* and *Fbp1^ΔHep^* mice (n=7).

(P) eWAT, SQ-WAT, BAT, kidney, and spleen weight/body weight ratio in 4 h fasted 8-wo *Fbp1^F/F^* and *Fbp1^ΔHep^* mice (n=7).

(Q) qRT-PCR of lipolytic mRNAs in eWAT of 4 h fasted 8-wo *Fbp1^F/F^* and *Fbp1^ΔHep^* mice (n=7).

(R) IB analysis of indicated proteins in eWAT of 4 h fasted 8-wo *Fbp1^F/F^* and *Fbp1^ΔHep^* mice (n=7). Densitometry determined relative normalized protein ratios (*Fbp1^ΔHep^/Fbp1^F/F^*) and P values are shown on the right.

(S-U) Serum lactate (S), insulin (T) and glucagon (U) in the indicated mice (n=7).

(V and W) Liver glycogen (V) and G6-P (W) in the indicated mice (n=7).

Data are presented as mean ± SEM. *P < 0.05, **P < 0.01 (Unpaired two-tailed t test). n.s., not significant.

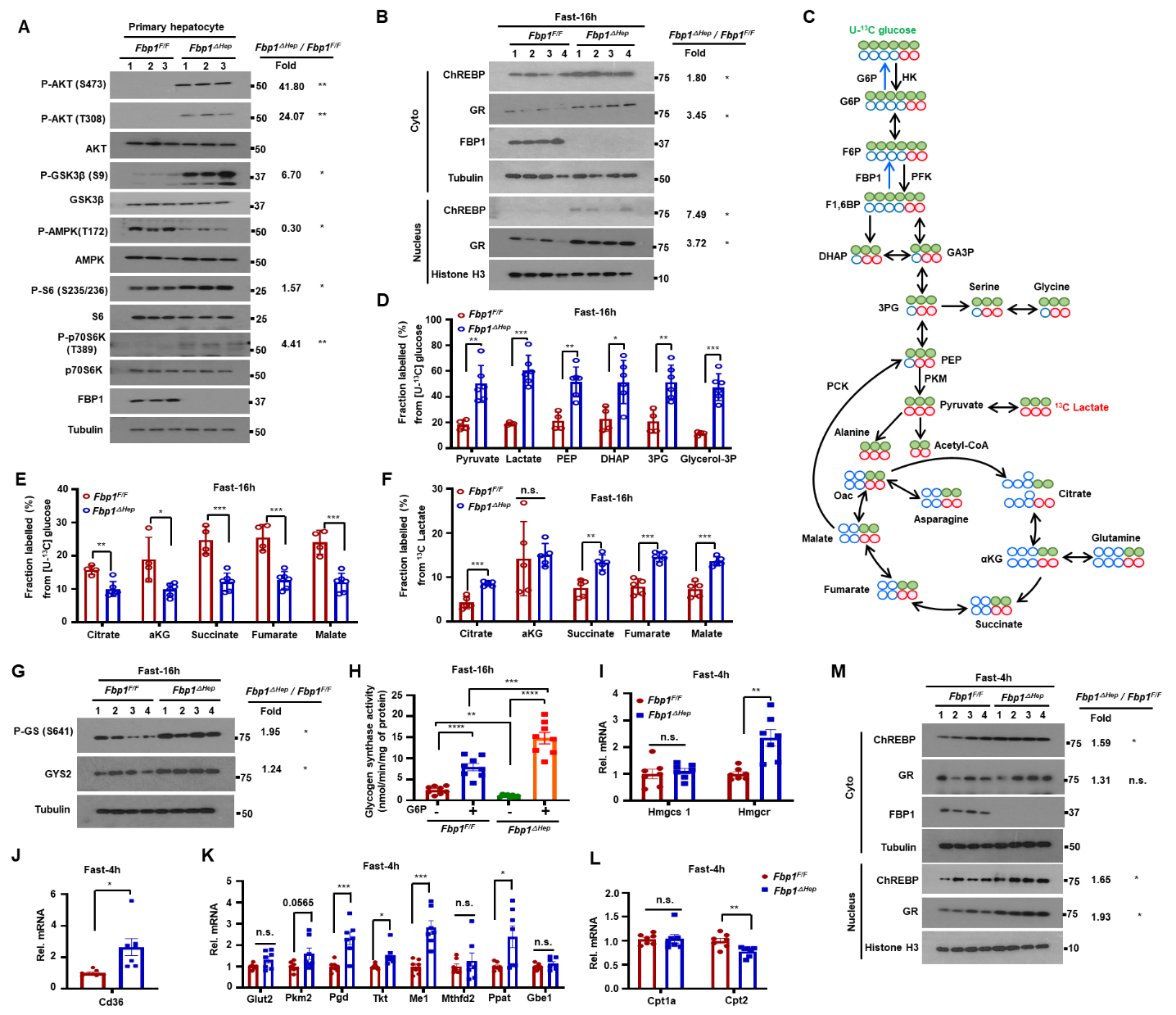

**Figure S2. Fasting induced metabolic genes and metabolic alterations in** ***Fbp1^ΔHep^*** **mice, related to Figure 2.**

(A) IB analysis of mouse primary hepatocytes from fasted 8-wo *Fbp1^F/F^* and *Fbp1^ΔHep^* mice. Relative normalized protein ratios (*Fbp1^ΔHep^/Fbp1^F/F^*) and P values are shown on the right.

(B) IB analysis of nuclear and cytosolic liver proteins in 16 h fasted 8-wo *Fbp1^F/F^* and *Fbp1^ΔHep^* mice. Relative normalized protein ratios (*Fbp1^ΔHep^/Fbp1^F/F^*) and P values are shown on the right.

(C) Schematic of U-^13^C glucose and ^13^C-Lactate tracing of 16 h fasted 8-wo *Fbp1^F/F^* and *Fbp1^ΔHep^* livers (n=4-6).

(D and E) Fractional labelling of glycolytic (D) and TCA (E) metabolites from U-^13^C glucose tracing of fasted 8-wo *Fbp1^F/F^* and *Fbp1^ΔHep^* liver (n=4-6).

(F) Fractional labelling of liver TCA intermediates from ^13^C-lactate tracing of indicated livers (n=5).

(G) P-GS(S641) and GYS2 in liver lysates of fasted 8-wo *Fbp1^F/F^* and *Fbp1^ΔHep^* mice. Relative normalized relative GYS2 and P-GYS2 ratios (*Fbp1^ΔHep^/Fbp1^F/F^*) and P values are shown on the right.

(H) Glycogen synthase activity in lysates of 16 h fasted 8-wo *Fbp1^F/F^* and *Fbp1^ΔHep^* livers (-/+) saturating exogenous G6-P (n=7-8).

(I-L) qRT-PCR of cholesterol synthesis (I), lipid uptake (J), glycolysis, pentose phosphate pathway, glycogen synthesis (K) and β-oxidation (L) related mRNAs in livers of 4 h fasted 8-wo *Fbp1^F/F^* and *Fbp1^ΔHep^* mice (n=7).

(M) IB analysis of nuclear and cytosolic extracts of 4 h fasted 8-wo *Fbp1^F/F^* and *Fbp1^ΔHep^* livers (n=7). Relative normalized protein ratios (*Fbp1^ΔHep^/Fbp1^F/F^*) and P values are shown on the right.

Data are presented as mean ± SEM, *P < 0.05, **P < 0.01, ***P < 0.001, ****p < 0.0001, n.s., not significant. (Unpaired two-tailed t test).

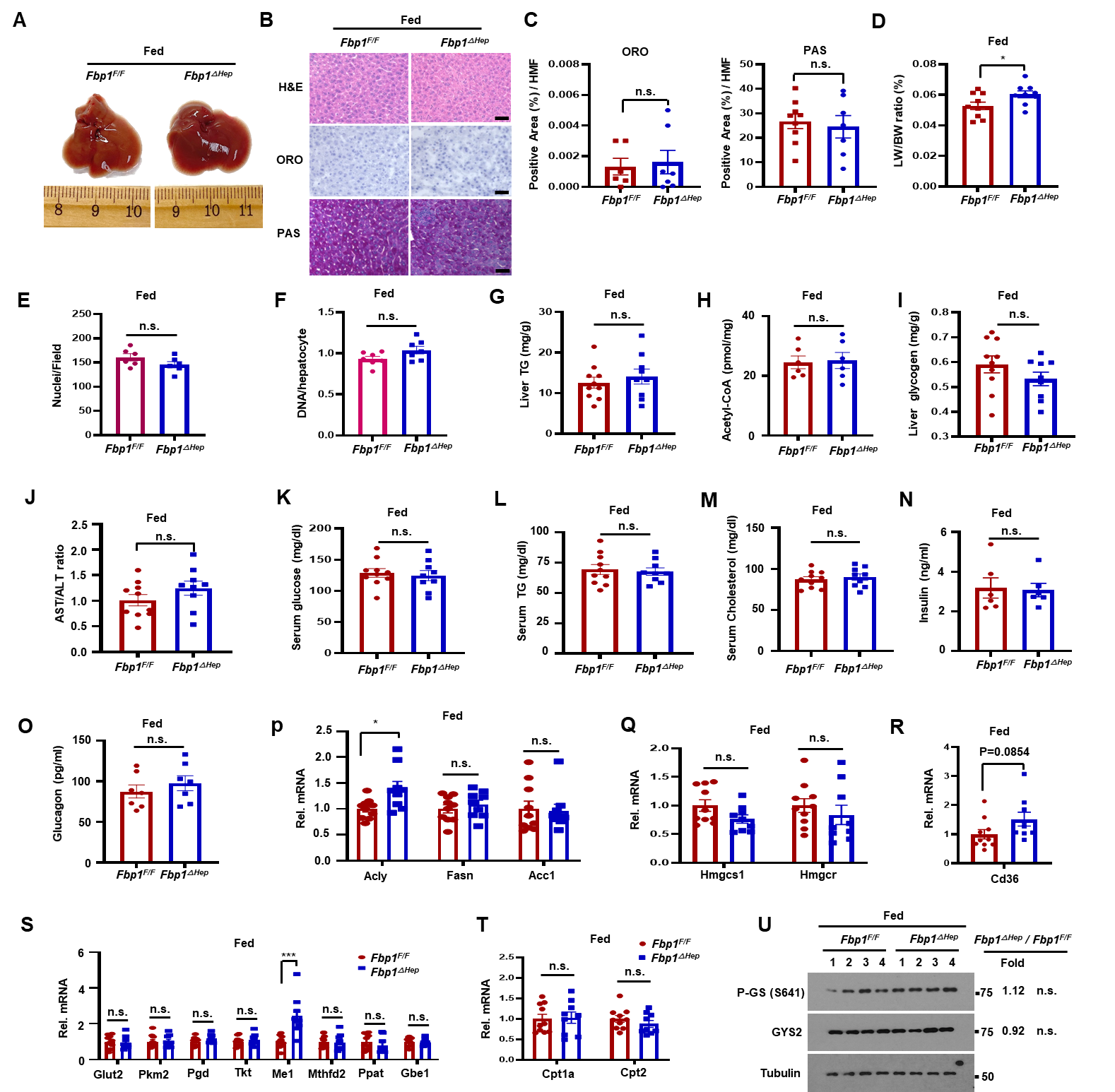

**Figure S3. The fed *Fbp1^ΔHep^* liver is metabolically normal, related to Figure 2.**

(A) Gross morphology of livers from 8-weeks-old (wo) *Fbp1^F/F^* and *Fbp1* *^ΔHep^* mice at the fed (*ad libitum*) state (n=9-10).

(B) H&E, ORO, and PAS staining of liver sections from the mice in S3A (n=9-10). Scale bars, 20 μm.

(C) ORO and PAS staining intensity per HMF were determined by Image J analysis of the liver sections in S3B.

(D-M) Liver/body weight (D), nuclei per field (E), hepatocyte DNA content (F), and liver TG (G), acetyl-CoA (H) and glycogen (I), and serum AST/ALT ratio (J), glucose (K), TG (L), and cholesterol (M) in the indicated mice (n=6-10).

(N and O) Serum insulin (N) and glucagon (O) in the indicated mice (n=6-10).

(P-T) qRT-PCR of lipogenesis (P), cholesterol synthesis (Q), lipid uptake (R), glycolysis, pentose phosphate pathway, glycogen synthesis (S) and β-oxidation (T) related mRNAs in livers of fed 8-wo *Fbp1^F/F^* and *Fbp1^ΔHep^* mice (n=9-10).

(U) P-GS(S641) and GYS2 in liver lysates of fed 8-wo *Fbp1^F/F^* and *Fbp1^ΔHep^* mice. Relative normalized protein ratios (*Fbp1^ΔHep^/Fbp1^F/F^*) and P values are shown on the right.

Data are presented as mean ± SEM, *P < 0.05, ***P < 0.001, n.s., not significant. (Unpaired two-tailed t test).

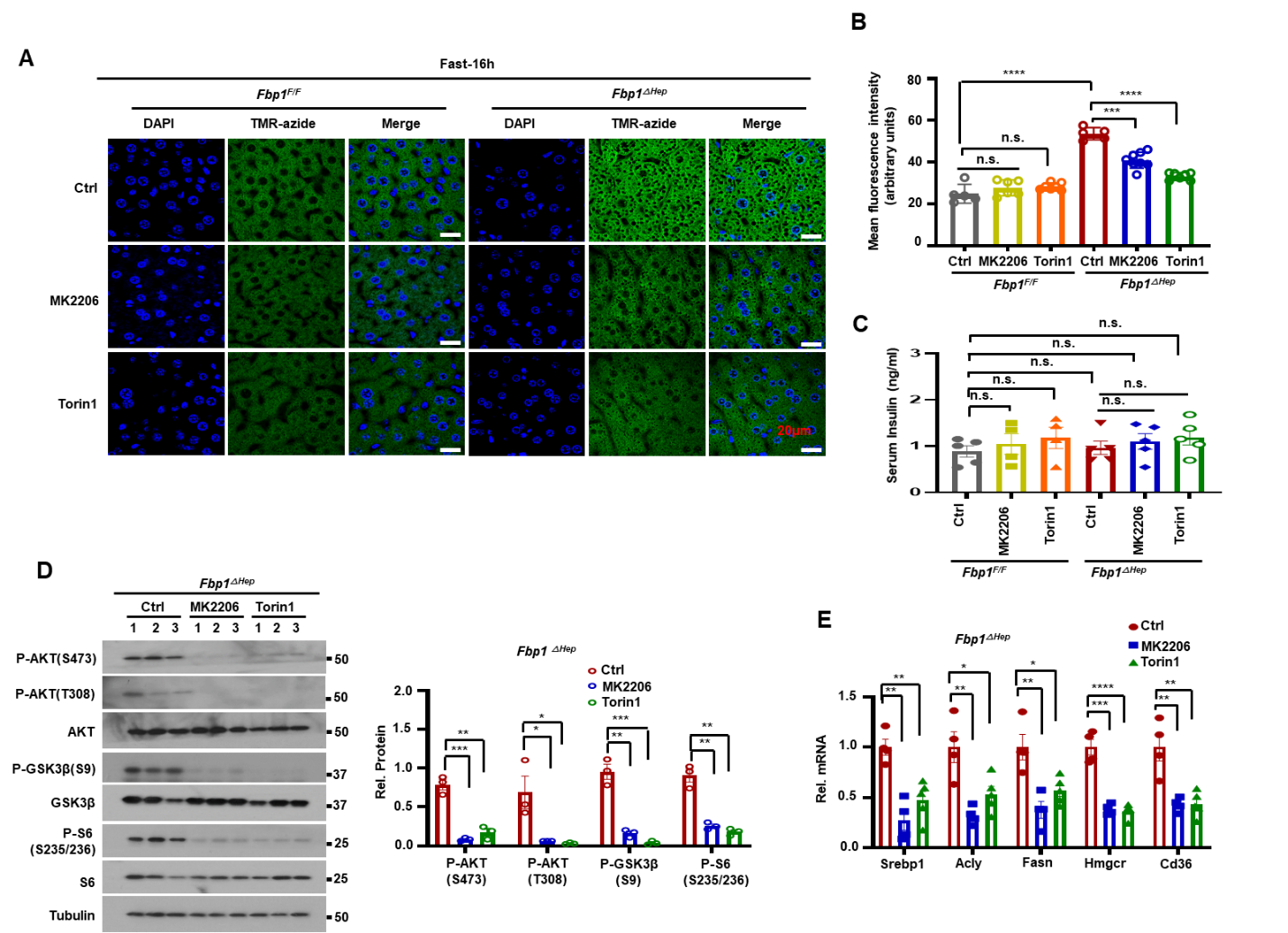

**Figure S4. AKT and mTORC inhibitors attenuate hepatomegaly, hepatosteatosis and hyperlipidemia in fasted *Fbp1^ΔHep^* mice, related to Figure 3.**

(A-B) Liver protein synthesis in untreated or MK2206 or Torin1 pretreated and 16 h fasted 8-wo *Fbp1^F/F^* and *Fbp1^ΔHep^* mice was determined by OP-puro (TMR-azide) incorporation (A) (n=4-5). Scale bars, 20 µm. Mean fluorescence intensity of above liver sections (B).

(C) Serum insulin levels in the mice from Figure 3A (n=4-5).

(D) IB analysis of liver lysates from above mice (left) and relative normalized protein amounts (right).

(E) qRT-PCR of liver *Srebp1c*, *Acly*, *Fasn*, *Hmgcr* and *Cd36* mRNAs in the mice from Figure 3A (n=4-5).

Data are presented as mean ± SEM. *P < 0.05, **P < 0.01, ***P < 0.001, ****p < 0.0001 (Unpaired two-tailed t test).

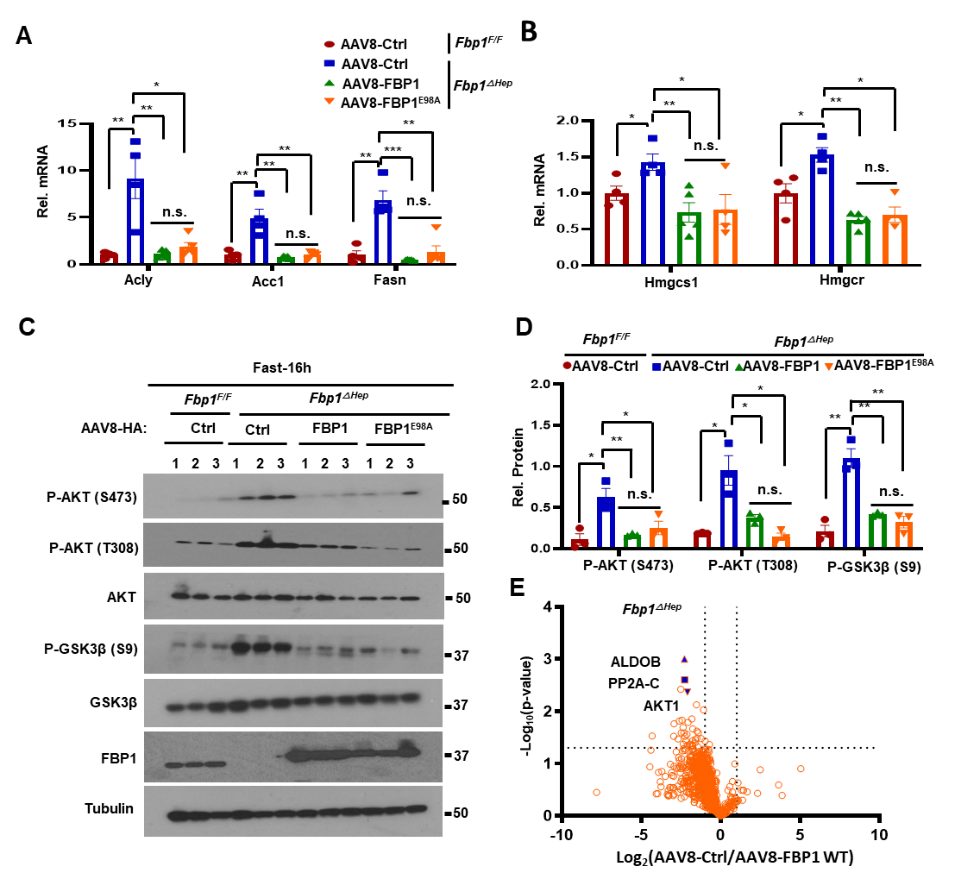

**Figure S5. Catalytically inactive FBP1 restrains hepatomegaly, fatty liver, and hyperlipidemia, but not hypoglycemia in *Fbp1^ΔHep^* mice, related to Figure 4.**

(A and B) qRT-PCR of liver *Acly*, *Acc1*, *Fasn* (A), *Hmgcs1* and *Hmgcr* (B) mRNAs of fasted *Fbp1^F/F^* and *Fbp1^ΔHep^* mice transduced with AAV8-Ctrl, AAV8-FBP1 and AAV8-FBP1^E98A^ (n=4-5).

(C and D) IB analysis of liver lysates from the mice in Figure 4A (C) and relative normalized protein amounts (D).

(E) Scatterplots of FBP1 interacting proteins identified by MS analysis of HA IPs from *Fbp1^ΔHep^* livers transduced with AAV8-HA-control or AAV8-HA-FBP1. The liver lysates were immuno-puriﬁed on anti-HA afﬁnity beads. The eluates were washed and analyzed by mass spectrometry.

Data are presented as mean ± SEM. *P < 0.05, **P < 0.01, ***P < 0.001, n.s., not significant. (Unpaired two-tailed t test).

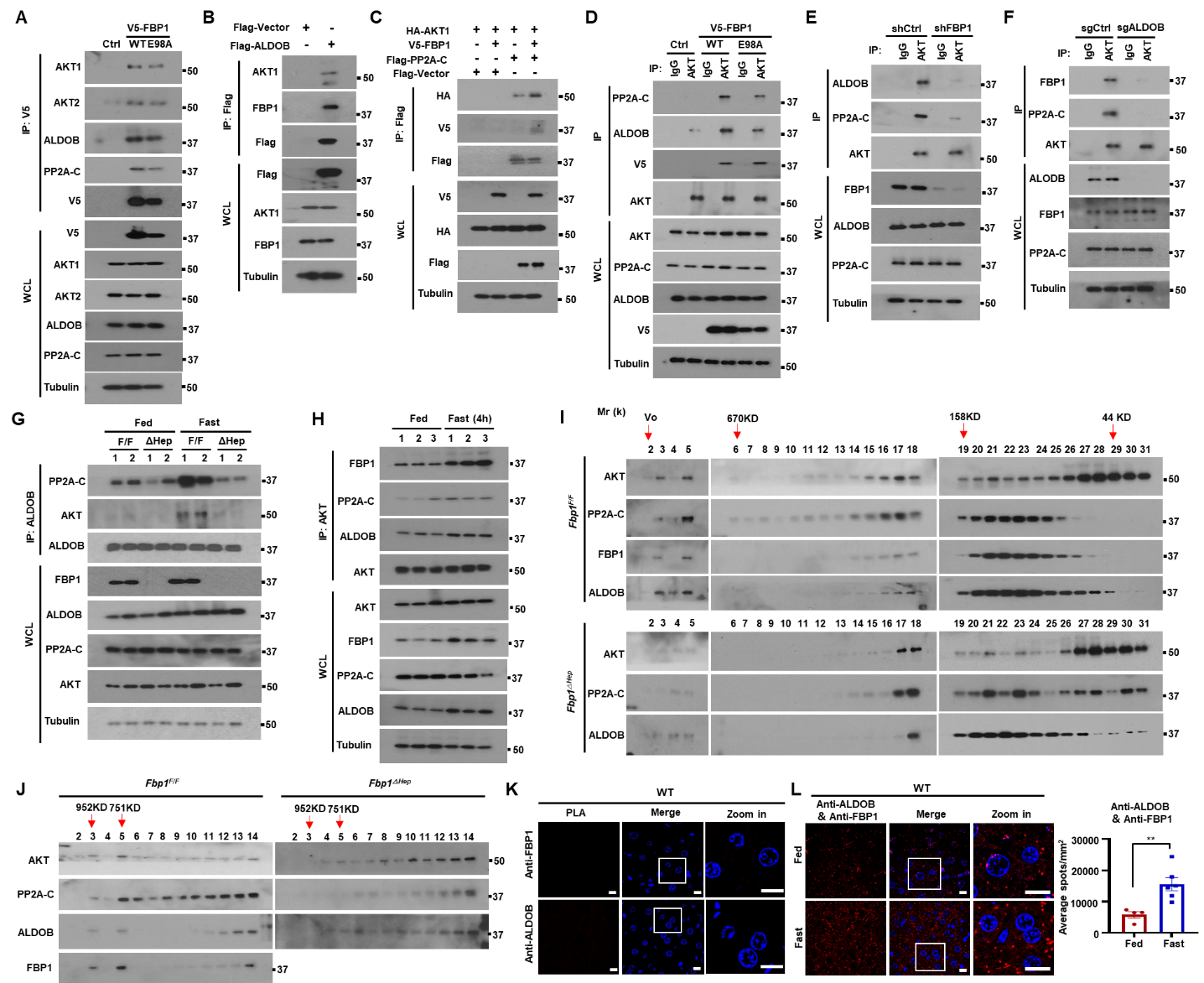

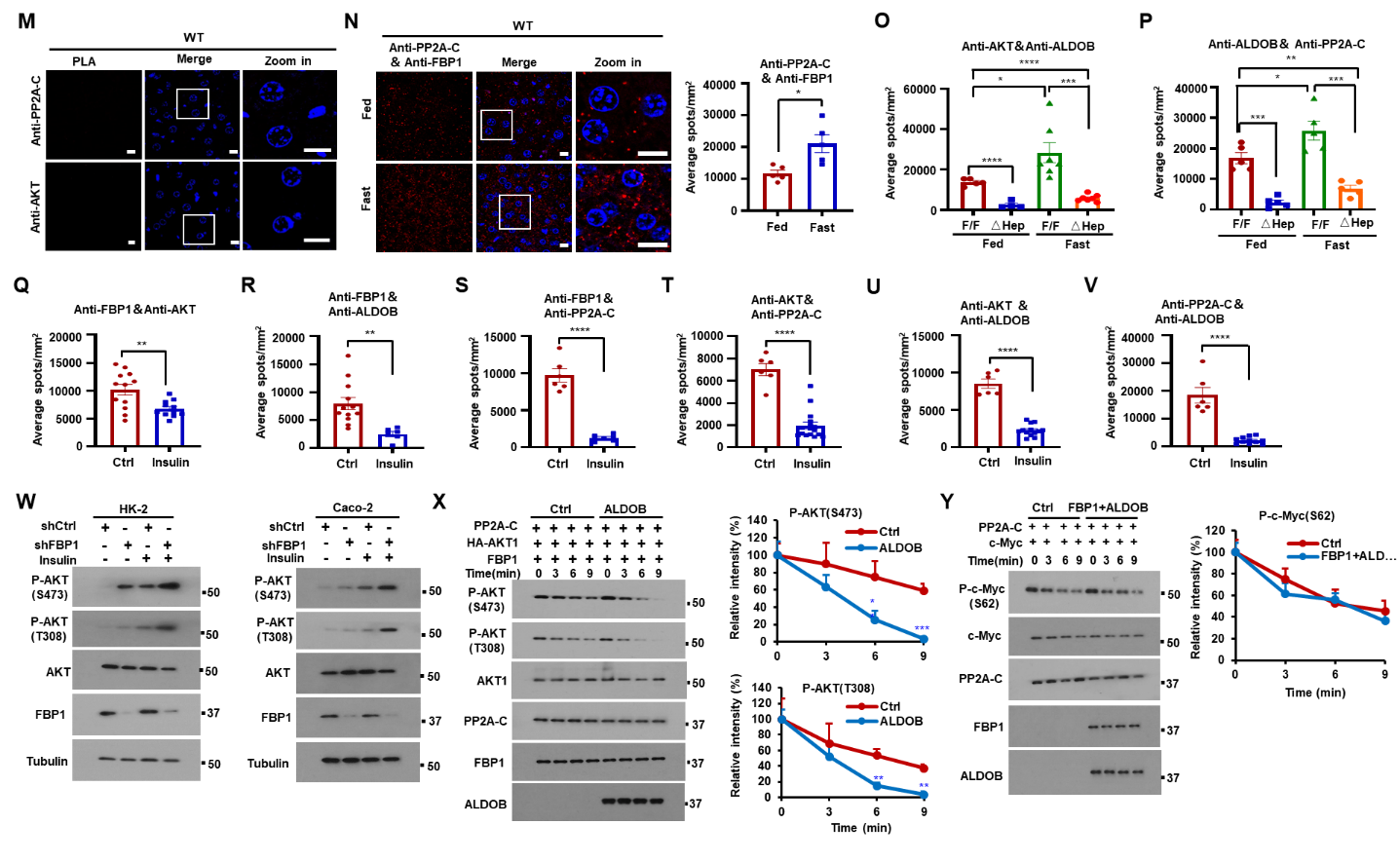

(Figure continued on next page)

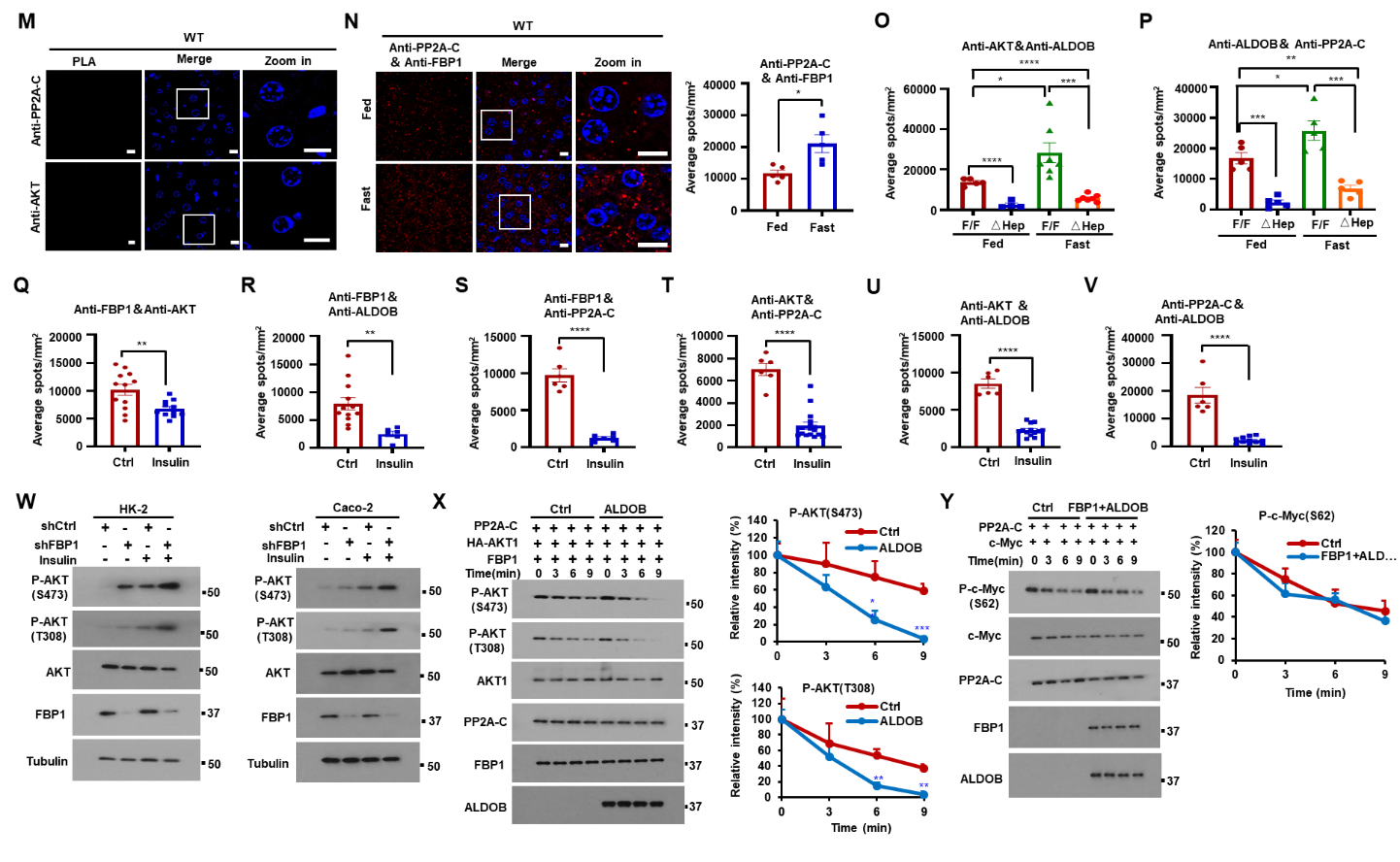

**Figure S6. FBP1 associates with ALDOB and PP2A-C to form a protein complex that binds AKT and inhibits its activation, related to Figure 5.**

(A) IP analysis of 293T cells transfected with Ctrl, V5-FBP1 or V5-FBP1^E98A^ vectors. The gel separated IPs were IB’ed with the indicated antibodies.

(B) IP analysis of 293T cells transfected with empty Flag or Flag-ALDOB vectors. The gel separated IPs were IB’ed with the indicated antibodies.

(C) IP analysis of 293T cells transfected with HA-AKT1, V5-FBP1 and Flag-PP2A-C. The gel separated IPs were IB’ed with the indicated antibodies.

(D) IP analysis of Huh7 cells stably transfected with FBP1, FBP1^E98A^ or Ctrl vectors. The gel separated IPs were IB‘ed with the indicated antibodies.

(E and F) IP analysis with indicated antibodies of Huh7 cells stably transfected with shCtrl, shFBP1 (E) and sgCtrl or sgALDOB (F). The gel separated IPs were IB‘ed with the indicated antibodies.

(G and H) IP analysis of liver lysates from 8-wo *Fbp1^F/F^* and *Fbp1^△Hep^* mice fed or fasted for 16 h (G) or 4h (H). The gel separated IPs were IB’ed with the indicated antibodies.

(I and J) Liver lysates of fasted 8-wo *Fbp1^F/F^* and *Fbp1^ΔHep^* mice were analyzed by gel filtration chromatography. Fractions were resolved by SDS-PAGE and probed with the indicated antibodies (n=3) (I). Fractions 2-14 were analyzed again as above (J). Arrows indicate the approximate elution volumes of molecular weight standards: blue dextran, Vo; thyroglobulin, 670kDa; γ-globulin, 158kDa; ovalbumin, 44kDa; and myoglobin, 17kDa.

(K) Representative images showing the absence of a PLA signal in liver sections of fasted 8-wo WT mice stained with either anti-FBP1 or anti-ALDOB alone followed by the PLA reaction (n=5). Scale bars, 10 μm.

(L) Representative images of ALDOB-FBP1 (left) interactions detected by PLA of liver sections of fed or 16 h fasted 8-wo WT and *Fbp1^ΔHep^* mice stained with anti-FBP1 and anti-ALDOB together (n=5). Scale bars, 10 μm. Quantification of representative PLA images is shown on the right.

(M) Representative images showing the absence of a PLA signal in liver sections of fasted 8-wo WT mice stained with either anti-PP2A-C or anti-AKT alone followed by the PLA reaction (n=5). Scale bars, 10 μm.

(N) Representative images of PP2A-C-FBP1 (left) interactions detected by PLA of liver sections of fed or 16 h fasted 8-wo WT mice stained with anti-FBP1 and anti-PP2A-C together (n=5). Scale bars, 10 μm. Quantification of representative PLA images is shown on the right.

(O and P) Quantification of representative images of ALDOB-AKT (O) and ALDOB-PP2A-C (P) interactions detected by PLA of liver sections of fed or 16 h fasted 8-wo WT and *Fbp1^ΔHep^* mice stained with the indicated antibody combinations (n=5).

(Q-V) Quantification of representative PLA images of AKT-FBP1 (Q), ALDOB-FBP1 (R), PP2A-C-FBP1 (S), AKT-PP2A-C (T), AKT-ALDOB (U), and ALDOB-PP2A-C (V) interactions detected by PLA of liver sections from fasted 8-wo WT mice treated -/+ insulin (0.5U/kg) stained with the indicated antibody combinations (n=5).

(W) HK-2 and CaCo-2 cells stably transfected with shCtrl or shFBP1 were treated -/+ insulin (100 nM) and IB’ed with indicated antibodies.

(X) In vitro dephosphorylation of HA-AKT1 isolated from insulin stimulated Huh7 cells and incubated with active-PP2A-C in the presence or absence of ALDOB. The reactions were IB analyzed with the indicated antibodies (left). Relative P-AKT (S473) and P-AKT (T308) intensity was determined by densitometry (right).

(Y) HA-c-Myc was isolated from EGF (100ng/ml) stimulated transiently transfected Huh7 cells incubated with active PP2A-C in the presence or absence of recombinant FBP1 and ALDOB. The reactions were IB analyzed with the indicated antibodies (left). Relative c-Myc (p-S62) intensity was determined by densitometry (right).

Data are presented as mean ± SEM. *P < 0.05, **P < 0.01, ***P < 0.001 ****p < 0.0001 (Unpaired two-tailed t test).

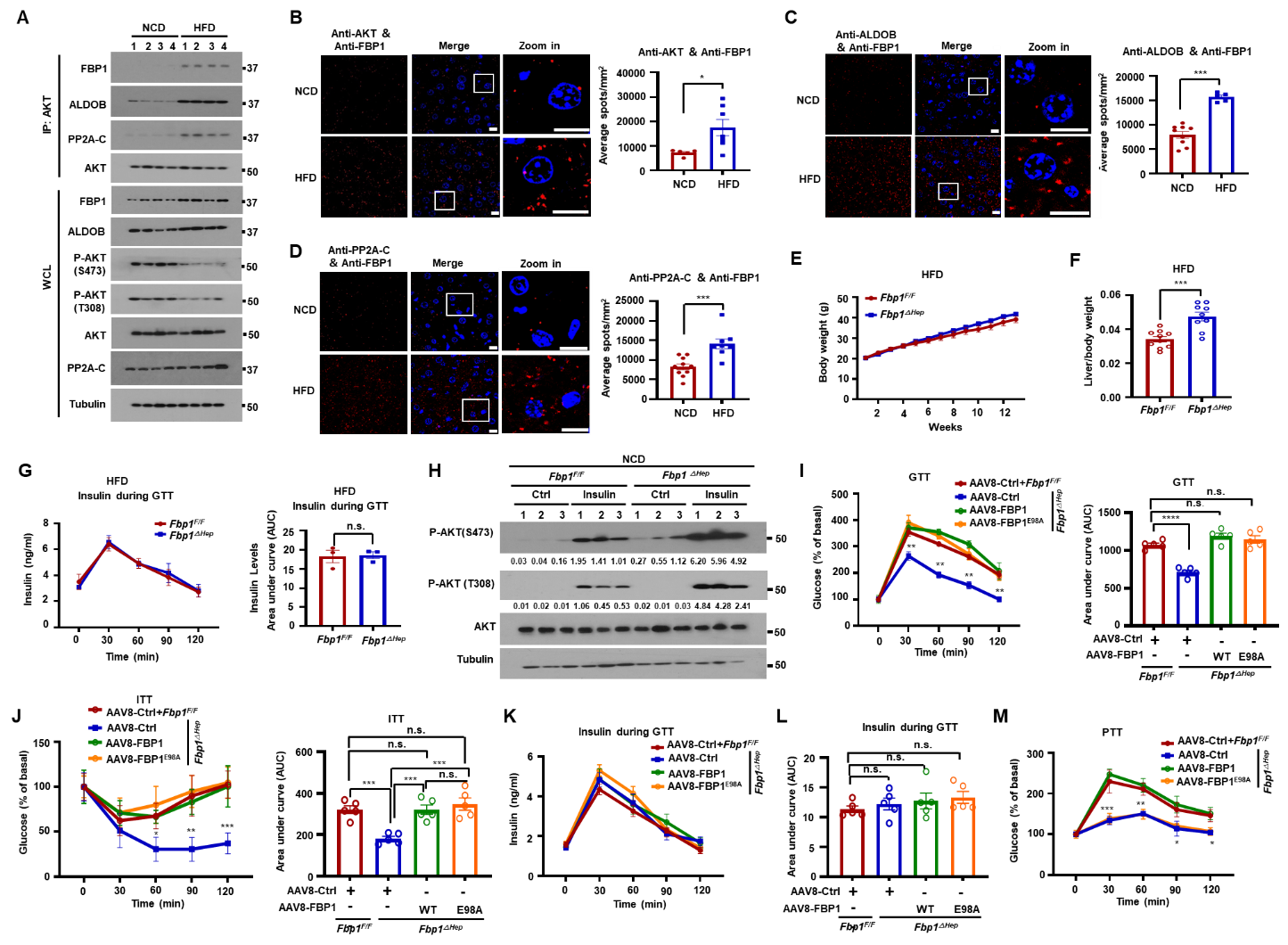

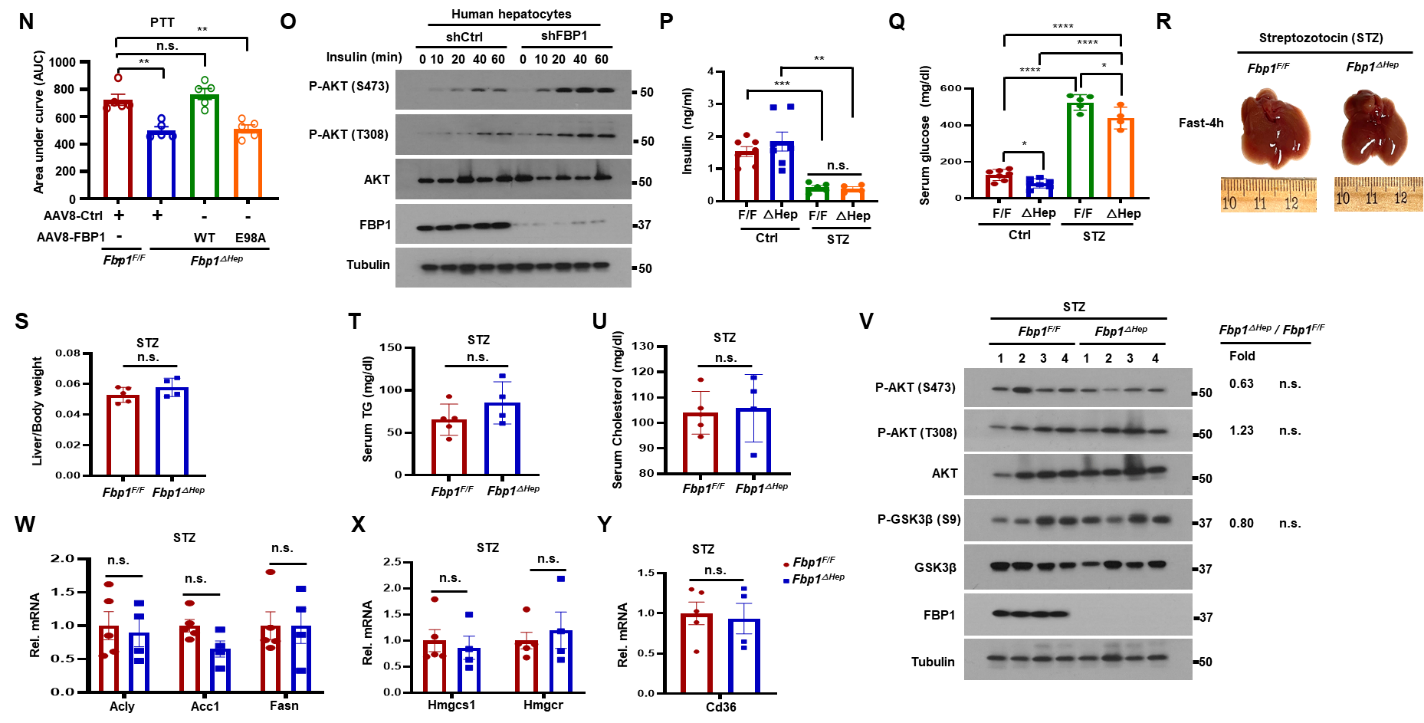

**Figure S7. *Fbp1^ΔHep^* mice are insulin hyperresponsive, related to Figure 6.**

(A) IP analysis of liver lysates from 18-wo NCD and HFD fed WT mice (n=4). The gel separated IPs were IB‘ed with the indicated antibodies.

(B-D) Representative images of AKT-FBP1 (B, left), ALDOB-FBP1 (C, left) and PP2A-C-FBP1 (D, left) interactions detected by PLA of liver sections of NCD or HFD fed WT mice stained with the indicated antibody combinations (n=5). Scale bars, 10 μm. Quantification of the PLA signals is shown on the right.

(E) Body weight gain by HFD-fed *Fbp1^F/F^* and *Fbp1^ΔHep^* mice (n=9-10).

(F) Liver/body weight ratio in indicated HFD-fed mice (n=9-10).

(G) Insulin levels during the GTT shown Figure 6I (n=3) (left) and AUC quantification (right).

(H) IB analysis of NCD-fed 8-wo *Fbp1^F/F^* and *Fbp1^ΔHep^* liver lysates prepared 15 min after control or insulin injections. Relative normalized protein amounts are shown below the P-AKT strips.

(I and J) GTT (I, left) and ITT (J, left) performed on fasted *Fbp1^F/F^* and *Fbp1^ΔHep^* mice transduced with AAV8-Ctrl, AAV8-FBP1 or AAV8-FBP1^E98A^ (n=4-5). AUC quantification of glucose amounts (I and J, right).

(K and L) Insulin concentrations during GTT of the indicated mice (K) (n=4-5) and AUC quantification (L).

(M and N) Pyruvate tolerance test (PTT) performed on fasted *Fbp1^F/F^* and *Fbp1^ΔHep^* mice transduced with AAV8-Ctrl, AAV8-FBP1 or AAV8-FBP1^E98A^ (n=4-5) (M). AUC quantification of glucose amounts (N).

(O) Human hepatocytes stably transfected with shCtrl or shFBP1 were stimulated with insulin (100 nM) for the indicated times after 6 h serum starvation. The cells were lysed and IB analyzed with the indicated antibodies.

(P and Q) Serum insulin (P) and glucose (Q) of -/+ STZ treated *Fbp1^F/F^* and *Fbp1^ΔHep^* mice (n=4-7).

(R) Gross liver morphology of 4 h fasted STZ treated 8-wo *Fbp1^F/F^* and *Fbp1^ΔHep^* mice (n=4-6).

(S-U) Liver/body weight ratio (S), serum TG (T) and cholesterol (U) of STZ treated *Fbp1^F/F^* and *Fbp1^ΔHep^* mice from S7R (n=4-6).

(V) IB analysis of liver lysates of STZ treated and fasted *Fbp1^F/F^* and *Fbp1^ΔHep^* mice. Relative normalized protein ratios (*Fbp1^ΔHep^/Fbp1^F/F^*) and P values are shown on the right.

(W-Y) qRT-PCR of *Acly*, *Acc1, Fasn* (W)*, Hmgcs1, Hmgcr* (X) and *Cd36* (Y) mRNAs in livers of STZ treated and fasted *Fbp1^F/F^* and *Fbp1^ΔHep^* mice (n=4-6).

Data are presented as mean ± SEM. *P < 0.05, **P < 0.01, ***P < 0.001, ****p < 0.0001, n.s., not significant (Unpaired two-tailed t test).

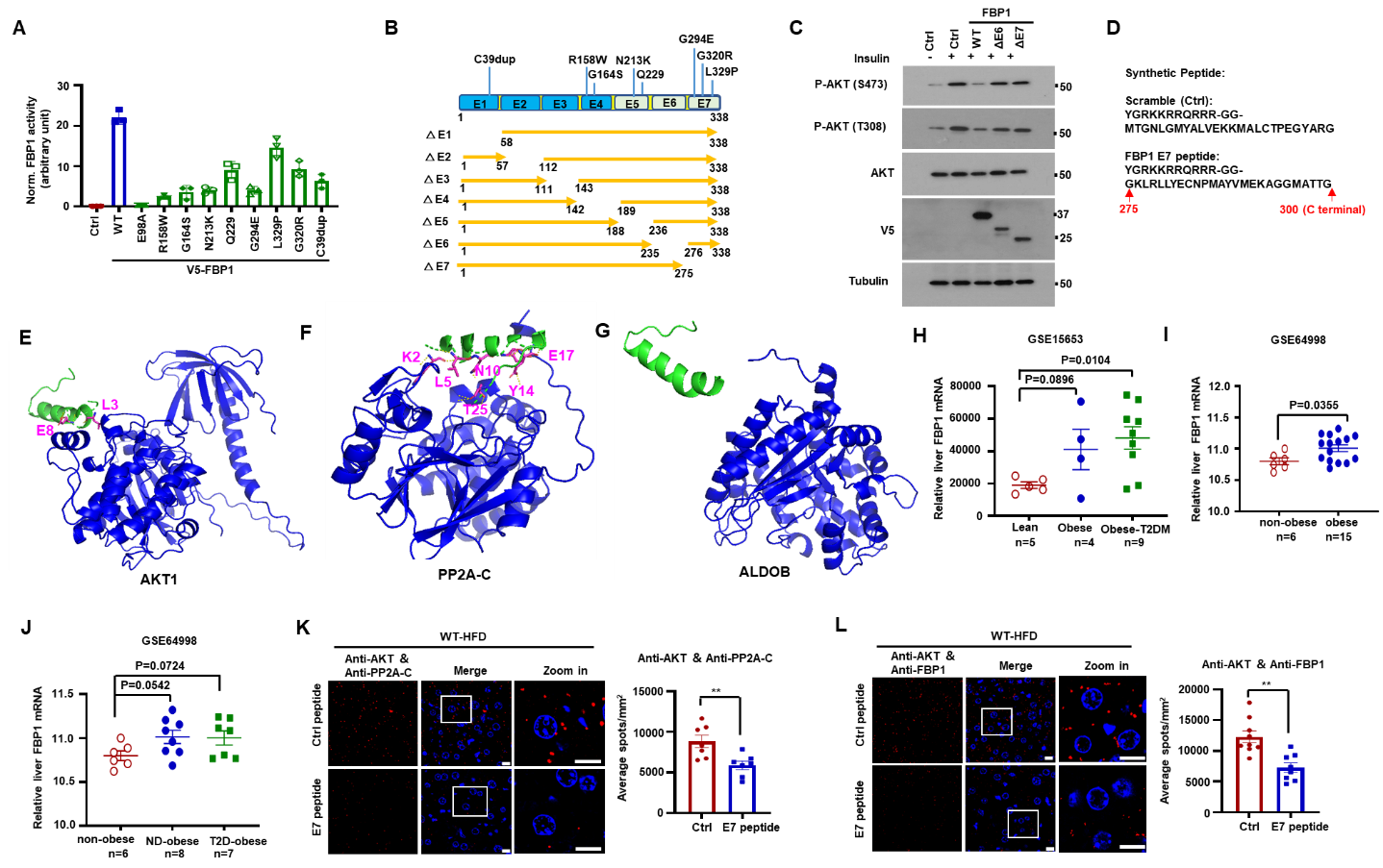

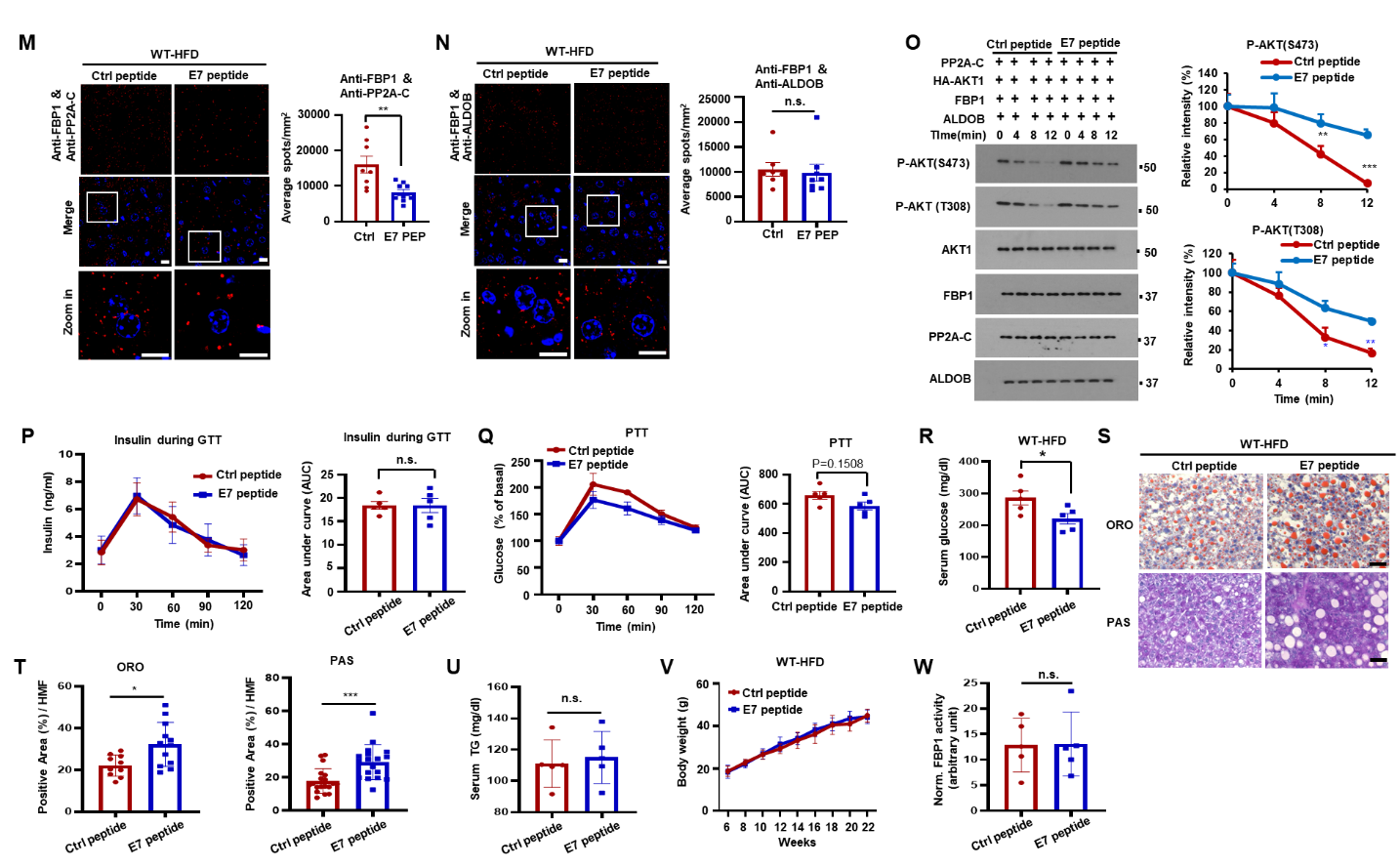

**Figure S8. An FBP1 C-terminal peptide that disrupts FBP1:AKT:PP2A-C interactions activates AKT and ameliorates insulin resistance, related to Figure 7.**

(A) Normalized FBP1 enzyme activities of WT and missense human FBP1 mutants expressed in Huh7 cells.

(B) Schematic of FBP1 exon deletion mutants.

(C) Huh7 cells stably expressing the indicated proteins were treated with -/+ insulin (100 nM) and IB’ed with the indicated antibodies.

(D) The sequence of the cell permeable E7 peptide (preceded by the TAT peptide) corresponding to FBP1 AA 275-300. The scrambled peptide was used as a control.

(E-G) AlphaFold predictions of E7 peptide docking onto AKT1 (E), PP2A-C (F) and ALDOB (G).

(H-J) Relative *FBP1* mRNA levels according to GEO databases GSE15653 of livers from lean, obese and obese T2DM (H), GSE64998 of livers from non-obese and obese individuals (I), GSE64998 of livers from non-obese, ND-obese and T2D-obese individuals (J). T2DM, [Type 2 diabetes. ND-obese, non-diabetic obese.](https://www.health.harvard.edu/a_to_z/type-2-diabetes-mellitus-a-to-z)

(K-N) Representative images of AKT-PP2A-C (K, left), AKT-FBP1 (L, left), FBP1-PP2A-C (M, left) and FBP1-ALDOB (N, left) interactions detected by PLA of liver sections of HFD-fed WT mice treated with Ctrl or FBP1 E7 peptides and stained with the indicated antibody combinations (n=5). Scale bars, 10μm. Quantification of representative PLA images is shown on the right.

(O) In vitro dephosphorylation of HA-AKT1 isolated from insulin stimulated Huh7 cells and incubated with active-PP2A-C in the presence or absence the E7 peptide. The reactions were IB analyzed with the indicated antibodies (left). Relative intensity of P-AKT (S473) and P-AKT (T308) was determined by densitometry (right).

(P) Insulin levels during the GTT from Figure 7H (left) and AUC quantification of the insulin levels (right) (n=5).

(Q) PTT performed on the HFD-fed mice from Figure 7D fasted for 12-14h (left) and AUC quantification (right).

(R) Serum glucose in mice from Figure 7D (n=5).

(S and T) ORO and PAS staining of liver sections from Figure 7D (S) (n=5). Scale bars, 20 μm. ORO and PAS staining positive areas per HMF (T).

(U) Serum TG in mice from Figure 7D (n=5).

(V) Body weight gain of HFD fed mice treated with Ctrl and FBP1 E7 peptide (n=5).

(W) Normalized FBP1 activity in HFD fed mice from Figure 7D (n=5).

Data are presented as mean ± SEM. *P < 0.05, **P < 0.01, ***P < 0.001, n.s., not significant (Unpaired two-tailed t test).

**Table S1. Primers for constructs used in this study.**

| **Insert** | **Forward Primer (5’-3’)** | **Reverse Primer (5’-3’)** |
| --- | --- | --- |
| FBP1-R158W | GCAACCAGGCTGGAACCTGGT | AGAGCATCCTTCTCAGAAGGC |
| FBP1-G164S | GGTGGCAGCCAGCTACGCACT | AGGTTCCGGCCTGGTTGC |
| FBP1-N213K | ACAGCCTTAAGGAGGGCTACG | AGATTTTACCTTTCTTTTTTATCTTCACATC |
| FBP1-Q229 | TGAGTACATCTAGAGGAAGAAGTTC | GTGACGGCAGGGTCAAAG |
| FBP1-G294E | AAGGCTGGGGAAATGGCCACC | CTCCATGACGTAGGCCATG |
| FBP1-G320R | GGTGATCTTGAGATCCCCCGAC | GGCGCCCTCTGGTGAATG |
| FBP1-L329P | CTCGAGTTCCCGAAGGTGTATGAGAAGC | CACGTCGTCGGGGGATCC |
| FBP1-C39dup | CACTGCACAGCAGTCAAAGCC | CAGGAGCGAGTTGAGCAGCTGG |

**Table S2. shRNA and sgRNA sequences used in this study.**

| **shRNA and sgRNA** | **Target Sequences（5’-3’）** |
| --- | --- |
| sgALDOB#1 | CACCGGATCACACCCCCGATGCTC |
| sgALDOB#2 | CACCGGATTTCTCGGAACTGCCGG |
| shPP2A-C#1 | TGGAACTTGACGATACTCTAA |
| shPP2A-C#2 | CCCATGTTGTTCTTTGTTATT |
| shFBP1#1 | CGACCTGGTTATGAACATGTT |
| shFBP1#2 | CAGCAGTCAAAGCCATCTCTT |

**Table S3. Quantitative PCR primers used in this study.**

| **Gene** | **Forward Primer (5’-3’)** | **Reverse Primer (5’-3’)** |
| --- | --- | --- |
| Acly | GCCAGCGGGAGCACATC | CTTTGCAGGTGCCACTTCATC |
| Acaca | TGACAGACTGATCGCAGAGAAAG | TGGAGAGCCCCACACACA |
| Fasn | GCTGCGGAAACTTCAGGAAAT | AGAGACGTGTCACTCCTGGACTT |
| Hmgcs1 | GCCGTGAACTGGGTCGAA | GCATATATAGCAATGTCTCCTGCAA |
| Hmgcr | CTTGTGGAATGCCTTGTGATTG | AGCCGAAGCAGCACATGAT |
| Tkt | CGAAACCCTCACAATGATCG | TTCCTCAGGTTCAGCAGCTC |
| Pgd | ATGCCAGGAGGGAACAAAG | GTTCTCCGGTTCCCACTTTT |
| Glut2 | TGCTGGCCTCAGCTTTATTC | TTTCTTTGCCCTGACTTCCTC |
| Pkm2 | GCTA TTCGAGGAACTCCGCC | AAGGTACAGGCACTACACGC |
| Me1 | GATGATAAGGTCTTCCTCACC | TTACTGGTTGACTTTGGTCTGT |
| Mthfd2 | ACAGATGGAGCTCACGAACG | TGCCAGCGGCAGATATTACA |
| Ppat | AATAGCTGTGGCCCATAACG | ACGTGGAAAGCCCAATACC |
| Gbe1 | GCGATCATGGAACATGCTTACTAT | CCATAACGACTTGAAGCTGCAA |
| Cpt1α | TGTCCAAGTATCTGGCAGTCG | CATAGCCGTCATCAGCAACC |
| Cpt2 | ATCGTACCCACCATGCACTAC | CTGTCATTCAAGAGAGGCTTCTG |
| Srebp1c | GGAGCCATGGATTGCACATT | GGCCCGGGAAGTCACTGT |
| Cd36 | TCCTCTGACATTTGCAGGTCTATC | AAAGGCATTGGCTGGAAGAA |
